# Cognitive Computational Model Reveals Repetition Bias in a Sequential Decision-Making Task

**DOI:** 10.1101/2024.05.30.596605

**Authors:** Eric Legler, Darío Cuevas Rivera, Sarah Schwöbel, Ben J. Wagner, Stefan Kiebel

## Abstract

Humans tend to repeat past actions due to rewarding outcomes. Recent computational models propose that the probability of selecting a specific action is also, in part, based on how often this action was selected before, independent of previous outcomes or reward. However, these new models so far lack empirical support. Here, we present evidence of a repetition bias using a novel sequential decision-making task and computational modeling to reveal the influence of choice frequency on human value-based choices. Specifically, we find that value-based decisions can be best explained by concurrent influence of both goal-directed reward seeking and a repetition bias. We also show that participants differ substantially in their repetition bias strength, and relate these measures to task performance. The new task enables a novel way to measure the influence of choice repetition on decision-making. These findings can serve as a basis for further experimental studies on the interplay between rewards and choice history in human decision-making.

## Introduction

Over a century ago, ***Thorndike (1911***) proposed the *law of effect*, which states that actions that lead to rewarding outcomes are more likely to be repeated. The law of effect gained widespread recognition and is considered an important foundation for the development of early operant conditioning (***Skinner, 1963***) and modern-day reinforcement learning (***Sutton and Barto, 2018***). What is less known is that Thorndike additionally stated the *law of exercise*, also known as the *law of use*, saying that humans tend to repeat previous actions regardless of reward (***Thorndike, 1911***).

Consider, for instance, the morning routine that many of us follow, e.g. we start with a cup of tea or coffee, take a shower, have breakfast, brush our teeth, and get ready for work. Although such action sequences may be learned only by goal-directed reward seeking (law of effect), such learning might also be based on Thorndike’s law of exercise. Indirect empirical evidence for the law of exercise, i.e. a measurable repetition bias, stems from questionnaire studies on everyday behavior (***Ouellette and Wood, 1998; Hagger et al., 2002; Wood et al., 2002; Neal et al., 2006; Verplanken, 2006; McCloskey and Johnson, 2019***), showing that participants reliably repeat behavior in a context-dependent manner, for example a specific morning routine or the mode of transportation to work.

Experimental evidence, across disciplines, shows that repetition affects human decision making and learning. It improves language learning (***Lynch and Maclean, 2000***), likely through increased word-familiarity (***Perea et al., 2016***) and better learning of multiword expressions (***Majuddin et al., 2021***). It also modulates learning of motor and cognitive skills (***Huang et al., 2011; Magallon et al., 2016; Reinkensmeyer et al., 2016; Wolpert et al., 2011; Spampinato and Celnik, 2021***) and affects memory retrieval, judgement (***Scarborough et al., 1977; Hintzman, 1976***) and working memory processes (***Oberauer et al., 2015***), demonstrating its broad influence on cognitive functions. Effects of repetition have also been studied under specific experimental conditions, suggesting their independence from direct reward. Here, repetition significantly affects perceptual decision making by accelerating response times for ambiguous stimuli (***Akaishi et al., 2014***). Similarly, repetition can alter preferences in value-based decision-making processes (***Nebe et al., 2024***), suggesting that the influence of repetition extends beyond the direct anticipation or receipt of reward, challenging standard views on the effect of reward on decision making and learning. Most importantly, repetitions are seen as a key element of habit formation (***Wood and Rünger, 2016; Watson et al., 2022***).

Over the last decade, the study of habitual vs. goal-directed responses have been enriched through a broad range of studies using devaluation tasks, extinction tests or more complex cognitive tasks, like the Wisconsin card-sorting task (***Wilson and Niv, 2012***) or the two-step task (***Daw et al., 2011***). These studies helped broaden our understanding of whether an action is outcome-oriented via a probabilistic map of the environment or rather driven by obtaining past reward in the same situation, as for example might be the reason for an insensitivity to devaluation. For instance, for the two-step task, behavior is described by using a mixture of model-based (MB) and model-free (MF) reinforcement learning (RL). Here, the MB controller learns a probabilistic action-outcome mapping, i.e. a more sophisticated goal-directed higher order cognitive process, while the MF controller is governed by simpler stimulus-response associations (***Daw et al., 2005, 2011***). This approach provided many insights on how humans learn and perform tasks (***Daw et al., 2011; Wunderlich et al., 2012; McDannald et al., 2012; Doll et al., 2015, 2016; Gläscher et al., 2010***), and also highlighted how impairments in model-based planning can be linked to psychiatric disease (***Gillan and Robbins, 2014; Gillan et al., 2016; Seow et al., 2021; Wyckmans et al., 2019; Voon et al., 2015***). However, it is still debated whether the reward-driven nature of model-free RL aligns with the concept of habits, which is not related to immediate reward, but to mere repetition of actions (***Wood and Rünger, 2016; Watson et al., 2022***). Two recent studies (***Miller et al., 2019; Schwöbel et al., 2021***) proposed a different mechanism. In these studies, based on simulations, behavior was explained by the interaction of two components. First, as usual, goal-directed behavior was explained by a model-based planner. Second, the novel proposal was to model the effect of a repetition bias, following Thorndike’s law of exercise, based on past choice counts alone, without regard to outcome or reward. This perspective is also related to minimizing the complexity of an action policy over time (***Gershman, 2020***).

Here, we followed this theoretical lead and assessed empirically, in human participants (*π* = 70), the effect of such a repetition bias on behavior. We used a computational model to disambiguate the effects due to repetition bias from effects due to goal-directed behavior driven by reward maximization. To capture both type of effects, we developed a novel Y-navigation task (Y-NAT) in which participants perform sequences of movements in a 5x5 grid world to collect a trial-specific number of points. The task was designed to fulfill the following three main objectives: First, trial-specific points establish a clear goal that will prompt goal-directed behavior in participants. Second, the combination of a relatively tight deadline and the requirement to plan four moves ahead (see Materials and Methods) challenges participants in their capacity to act in a purely goal-directed fashion. Third, participants were informed about a so-called default action sequence (DAS), providing them with a less complex go-to strategy, which induces repetition of the same sequence over trials. The Y-NAT therefore enabled us to test (1) whether a repetition bias develops over time, (2) what its effects are on task behavior and (3) what the link is between individual differences in repetition bias and overall task-performance. Data was analyzed using both standard behavioral analyses and Bayesian model-based analyses. We used Bayesian model comparison to test several alternative models, with or without repetition bias.

## Results

We created a sequential decision-making task, the Y-navigation task (Y-NAT), to show the repetition bias (see Materials and Methods). For this grid-world task, participants had to collect points with four moves within a time limit of 6s and match a trial-specific goal sum of points as closely as possible (see ***Figure 1***). Participants were required to complete 16 blocks, each comprising 20 trials, resulting in a total of 320 trials.

**Figure 1.**
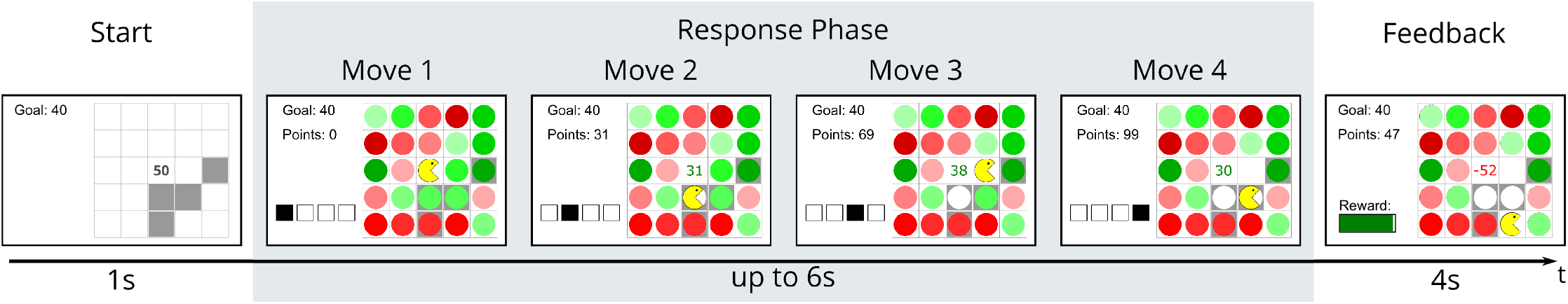
Illustration of a single task trial. At trial start, the goal, the fields of the DAS and the mean points of the DAS were shown on the screen for one second. During the subsequent response phase of up to six seconds, four moves had to be performed. During the feedback phase of 4 seconds at the end of each trial the reward was communicated.

In order to ensure the frequent repetition of at least one sequence of actions, a default action sequence (DAS) was highlighted. Using the DAS resulted in the highest expected reward in less than half of the trials (43.75%), with the lowest expected reward of the DAS being about half of the maximum reward. Furthermore, in half of the blocks, a probabilistic bonus could be earned when using the DAS.

In what follows we first present the results from standard behavioral analyses based on summary statistics and then move on to a model-based analysis.

### Behavioral Analysis

Our first approach was to find evidence of a repetition bias using inference statistics. For our task, we expected that a repetition bias manifests in the following ways: (1) an increase, over the course of the experiment, in the usage of the most frequently used sequence of actions; (2) an increase, over the course of the experiment, in the selection of the most frequently used sequence of actions in trials where this sequence of actions did not have the highest expected reward; (3) an increase, over the course of the experiment, to perform parts of the most frequently used sequence of actions, and (4) a decrease, over the course of the experiment, in the number of different sequences of actions being used.

We determined the proportion of trials in which the default action sequence (DAS) was executed, *p*(DAS), for each participant. As expected, the DAS was used in more than half of the trials (*p*(DAS) = .54, *SD* = .19), and 66 participants (94%) used the proposed DAS most frequently (see ***Table 1***). We found the expected difference in the proportion of DAS choices between the bonus (*p*(DAS) = .57, *SD* = .19) and the no bonus condition (*p*(DAS) = .52, *SD* = .19), *p <* .001, *d* = .26 (see Appendix ***Table 1*** for all descriptive statistics depending on the bonus condition).

**Table 1.**
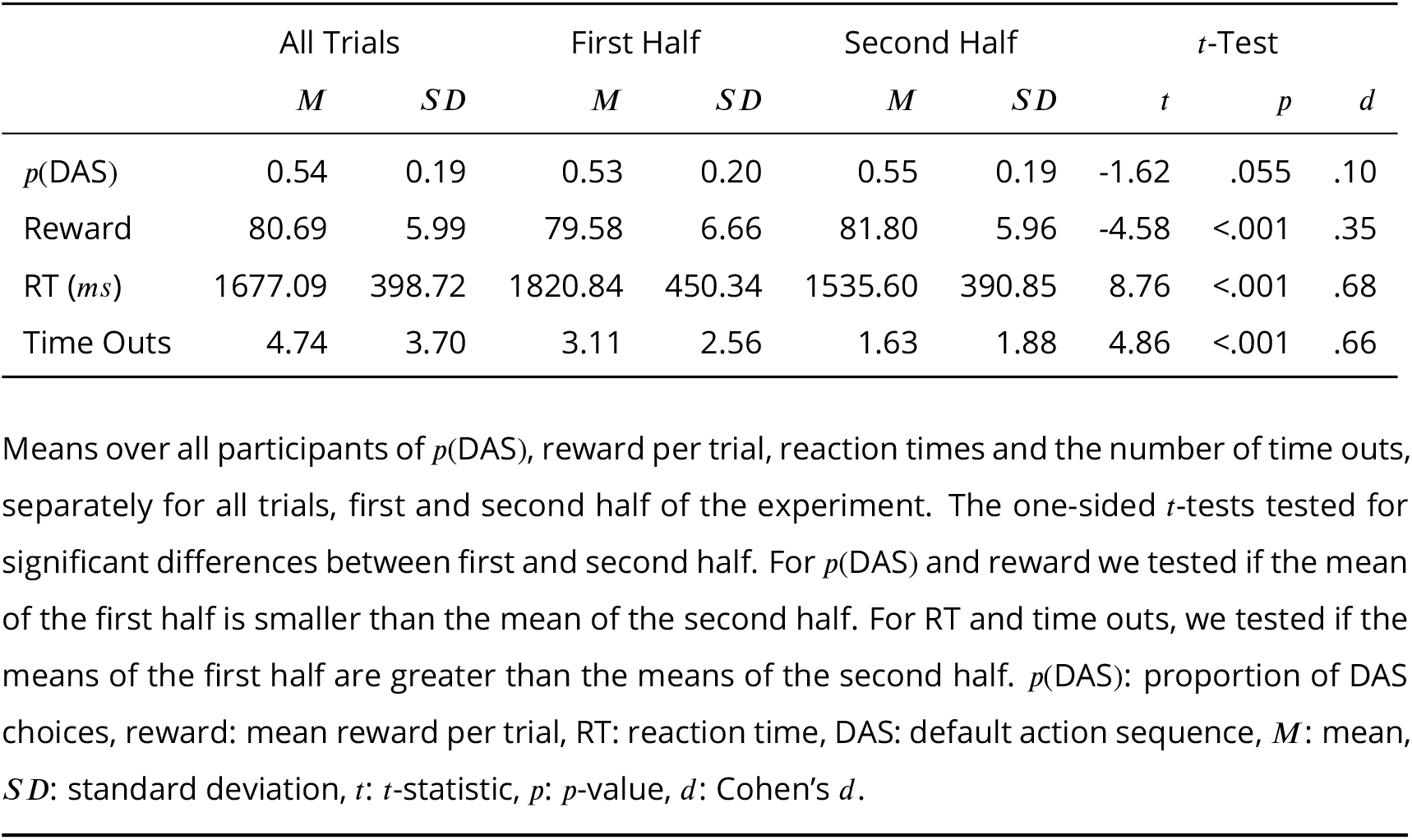
Descriptive statistics of performance measures for all trials and halves of the experiment

|  | All Trials |  | First Half |  | Second Half |  | *t*-Test |  |  |
| --- | --- | --- | --- | --- | --- | --- | --- | --- | --- |
|  | *M* | *SD* | *M* | *SD* | *M* | *SD* | *t* | *p* | *d* |
| *p*(DAS) | 0.54 | 0.19 | 0.53 | 0.20 | 0.55 | 0.19 | -1.62 | .055 | .10 |
| Reward | 80.69 | 5.99 | 79.58 | 6.66 | 81.80 | 5.96 | -4.58 | <.001 | .35 |
| RT (*ms*) | 1677.09 | 398.72 | 1820.84 | 450.34 | 1535.60 | 390.85 | 8.76 | <.001 | .68 |
| Time Outs | 4.74 | 3.70 | 3.11 | 2.56 | 1.63 | 1.88 | 4.86 | <.001 | .66 |

Means over all participants of *p*(DAS), reward per trial, reaction times and the number of time outs, separately for all trials, first and second half of the experiment. The one-sided *t*-tests tested for significant differences between first and second half. For *p*(DAS) and reward we tested if the mean of the first half is smaller than the mean of the second half. For RT and time outs, we tested if the means of the first half are greater than the means of the second half. *p*(DAS): proportion of DAS choices, reward: mean reward per trial, RT: reaction time, DAS: default action sequence, *M*: mean, *SD*: standard deviation, *t*: *t*-statistic, *p*: *p*-value, *d*: Cohen's *d*.

We focused all subsequent analyses on the DAS because participants used the DAS more frequently than expected when considering expected rewards, and the DAS was used most frequently by nearly all of the participants. We conducted three statistical analyses to test for a repetition bias: we tested for an increased usage of this sequence over time, an increased usage of this sequence even when the sequence did not yield the highest expected reward over time, and an increased usage of parts of the sequence, when the full sequence was not performed. Furthermore, we tested for a decrease of behavioral variability as an indicator of a repetition bias. We describe the results of these analyses in the following sections.

### Increase of DAS usage

We assessed whether there was an increase in DAS usage throughout the experiment. We repeated the differences between the trial-specific goals and the expected points of the DAS across halves and four segments (see ***Figure 8***D), and consequently expected rewards for the DAS to be repeated across the halves and the segments. The expected points for the DAS were communicated at the beginning of each trial and participants were able to calculate the expected reward for the DAS. Therefore, participants’ proportion of DAS choices should not change if they were guided only by expected rewards. However, if a repetition bias influenced participants’ choices, the DAS usage should have increased with time.

We compared the average proportion of DAS choices of all participants between the first and second half of the experiment, and over the four segments (comprising four subsequent blocks, see also ***Figure 8***D in Materials and Methods). During the first half, over all participants, the DAS was used in 53.3% (*SD* = 19.7%) of the trials, whereas in the second half the DAS was used in 55.2% (*SD* = 18.8) of the trials (see ***Figure 2***A). A one-sided *t*-test for related samples based on the individual differences of all participants only showed a non-significant difference, *t*(69) = −1.62, *p* = .055, *d* = .10. Similarly, there was only a non-significant increase of DAS usage across the four segments, repeated measures ANOVA with *F* (3, 207) = 1.52, *p* = .21,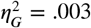, see ***Figure 2***A.

**Figure 2.**
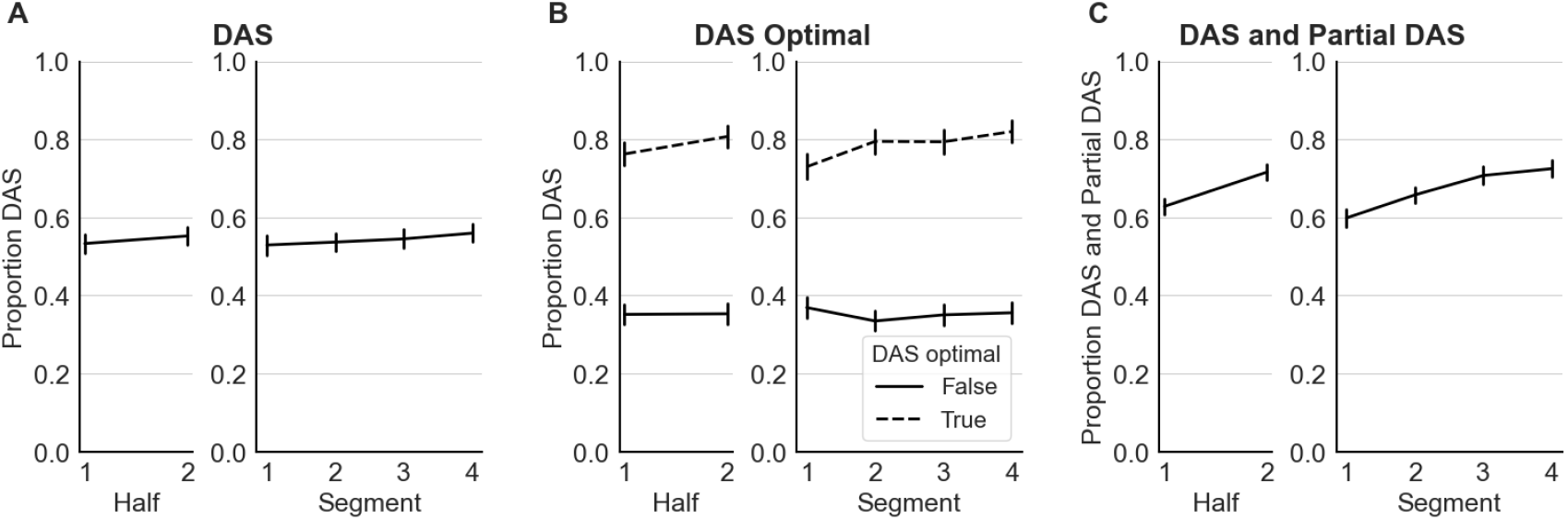
Default action sequence (DAS) choice behavior. **(A)** Mean proportional use of default action sequence (DAS) over the two halves (early/late) and four segments of the experiment. **(B)** Mean proportional use of default action sequence (DAS) depending on if the DAS was one of the sequences with the highest expected reward (optimal), separated by halves and by the four segments. **(C)** Mean proportional use of partial DAS use for non-DAS trials depending on halves and the four segments of the experiment. Black lines represent means over all participants. Error bars represent standard errors (*SE*).

### Increase in DAS usage in trials where DAS is not optimal

Although we did not find a significant increase in DAS usage over the course of the experiment, a repetition bias for the DAS should increase the probability of selecting the DAS irrespective of the expected reward of the DAS. We expected this because at the beginning of each block the DAS was an optimal choice (see Materials and Methods and ***Figure 8***B) participants were incentivized to use the DAS in the first trials of each block; this incentivized repetition of the DAS would bias participants towards choosing the DAS even when it did not yield the highest expected reward. However, the repetition bias could also decrease the probability to switch back to the DAS when the DAS is optimal later during the block. These two opposing effects together could explain why we found no significant overall increase in DAS usage.

To find this effect, we split up trials based on whether the DAS was one of the available sequences of actions with the highest expected reward, or not. We determined a use of DAS as optimal if the absolute difference between the points obtained by the DAS and the goal was ::; 5 points, because the difference between two adjacent colors was 10 points (see Materials and Methods). We conducted a repeated measures ANOVA with the proportion of DAS choices as dependent variable and the factors (1) halves of the experiment, and (2) DAS optimality. The main effect of expected reward was significant, *F* (1, 69) = 293.89, *p <* .001,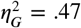. The main effect experimental halves was not significant, *F* (1, 69) = 3.88, *p* = .053,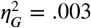(see ***Figure 2***B). We repeated this analysis with four segments instead of halves of the experiment as a factor. Again, the main effect of expected reward was significant, *F* (1, 69) = 232.15, 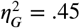 and the main effect of segment was not significant, *F* (3, 207) = 2.36, *p* = .077, 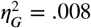 (see ***Figure 2***B).

Taken together, we did not find evidence for an increase in DAS use for trials where the DAS was not one of the sequences of actions with the highest expected reward. Hence, the potential repetition bias, established through the repeated use of the DAS in trials where the DAS is one of the sequences of actions with the highest reward, did not lead to an increase of DAS use at trials where the DAS did not have the highest expected reward.

### Increase in DAS parts

As participants had to execute a sequence of four moves in each trial, a repetition bias may have expressed itself by an increase of the probability of repeating at least the first move(s) of a sequence of actions. Due to the small trial-wise changes of the goals (see ***Figure 8***B), a possible strategy for participants would be to repeat the first move(s) of a sequence of actions but deviate from this sequence after these initial move(s), depending on the goal points.

We categorized the used sequences of actions into three categories: a DAS trial (when the full DAS was executed), a partial DAS trial (trial with a deviation from the DAS), or a DAS-independent trial (a completely different sequence). Trials that were categorized as partial DAS trials were defined by selecting at least the first move in accordance with the DAS, but not using the complete DAS.

To test for an increase in repeating the first move(s) of the DAS or the complete DAS, we compared the proportion of combined DAS and partial DAS trials to DAS independent trials, again with factor halves or segments. A one-sided *t*-test for related samples revealed that the proportion of combined DAS and partial DAS trials significantly increased from the first half of the experiment to the second half, *t*(69) = −6.46, *p <* .0001, *d* = .49 (see ***Figure 2***C). Similarly, a repeated measures ANOVA over segments showed a significant increase of combined DAS and partial DAS use over time, *F* (3, 207) = 19.53, *p <* .001, 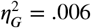 (see ***Figure 2***C).

### Decrease in behavioral variability

A repetition bias might also lead to a higher probability of repeating performed sequences of actions other than the DAS. This would lead to a lower number of different action sequences being performed in later stages of the experiment. To test this, we analyzed the number of used different sequences of actions between the halves and the four segments of the experiment. The mean number of used sequences of actions showed a small significant decrease from the first (16.87, *SD* = 5.94) to the second half (15.30, *SD* = 5.58), *t*(69) = 3.53, *p <* .001, *d* = .27 (see ***Figure 3***A). A repeated measures ANOVA with segments as factor showed a significant effect, *F* (3, 207) = 13.88, 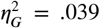(see ***Figure 3***B). Here the number of used sequences decreased significantly from the first to the second segment, but was stable throughout the following segments.

**Figure 3.**
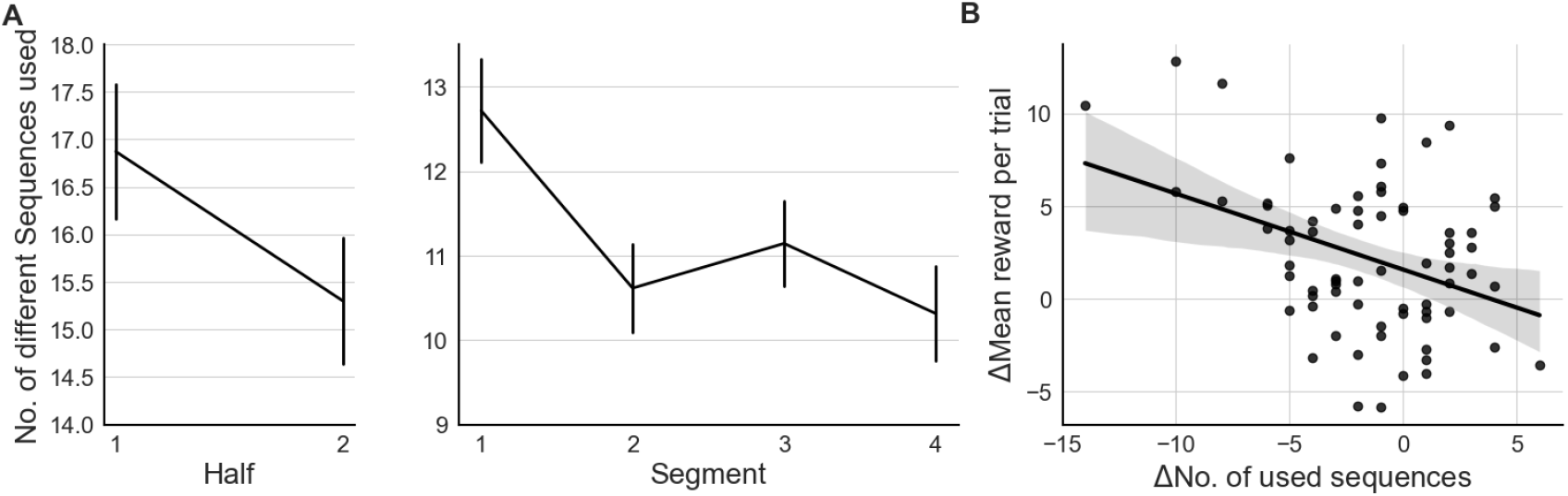
Decreasing behavioral variability of different sequences used. **(A)** Mean number of used different sequences of actions over the two halves (early/late) and four segments of the experiment. Error bars represent standard errors (*SE*). **(B)** Correlation between the difference of used sequences and the difference of mean reward per trial between the first and the second half. Each point represents one participant. The thick solid line represents linear regression model fitted to the data.

A decrease in behavioral variability could also reflect learning. Maybe participants selected some sequences of actions with very low rewards once during the first half of the experiment, but learned how to avoid selecting sequences with low rewards. To assess if this decrease in behavioral variability was caused by learning to select rewarding sequences more easily, we calculated the correlation between the difference of used sequences and the difference of mean reward per trial between the first and the second half. A negative correlation would indicate that participants who performed fewer sequences improved their ability to find rewarding sequences. This correlation showed a significant relationship in the expected direction, *r* = −.38, *p* = .001 (see ***Figure 3***C). We interpret this as evidence that the decrease in behavioral variability was probably caused by learning to select more rewarding sequences, over the course of the experiment.

In summary, we found only weak evidence for a repetition bias with summary statistics. Participants used the DAS more frequently than expected, but we only found hints for a repetition bias when also considering partial DAS choices or by analyzing the development of the number of used sequences. Conducting a similar analysis on the second most used sequence of actions or a Chi-square test of independence with all sequences of actions would not be meaningful: In contrast to the DAS, the expected rewards for all other sequences of actions differed between the first and second half of the experiment due to the grid layout changes between blocks. Hence, a sequence of actions that was frequently used during the first half of the experiment may generate very low rewards in the second half. As the repetition bias only increases the probability to repeat actions, we expect that action selection is mostly still guided by expected rewards, and participants should prefer to switch to action sequences with higher rewards (***de Wit et al., 2018; Watson and De Wit, 2018***). Therefore, to test for a repetition bias, the analysis should also take into account expected rewards. For this reason, we next turn to a model-based analysis, in which we consider simultaneously the repetition of action sequences and the expected rewards as effects on observed choices.

### Model-Based Analysis

A potential issue with our analyses above is the limited focus on behavioural measures for one specific sequence of actions, e.g. how many times the DAS was used in the first and second half of the experiment, thereby not considering expected rewards and other action sequences.

To consider all sequences of actions, and expected reward and repetition bias simultaneously, we used an adapted version of the prior-based control model (***Schwöbel et al., 2021***), which we call here the ‘expected value with proxy reward and repetition bias model’ (EVPRM). For the full model specification and details, see Materials and Methods and ***Figure 9***.

This model calculates the probability of selecting an action based on the balance of two components: the probability to repeat actions, and expected rewards. Crucially, the influence of repeated behavior is modeled by counting the number of times each action sequence has been used in the past *γ* _*π*_. This component is weighted by a free model parameter *α* _init_ that determines the strength of the repetition bias (see Expected value with proxy and repetition bias model (EVPRM)). Using this model, the focus is not restricted to one sequence of actions and the repetition bias parameter quantifies the influence of past behavior on action selection for all possible sequences of actions.

Moreover, the EVPRM incorporates the influence of expected rewards of all sequences of ac-tions on action selection. Like past behavior, expected rewards are weighted by a free parameter *β* that quantifies the individual precision on expected rewards. A high precision leads to pronounced probabilities and a stronger influence of expected rewards on action selection, while a low precision leads to more uniformly distributed probabilities and lower influence of expected rewards on action selection.

As the influence of expected reward and past behavior is modeled by two different parameters, *α* _init_ and *β*, we can disambiguate between effects on behavior by a low precision on expected rewards and effects on behavior driven by a strong repetition bias. Crucially, as we will show below this makes it possible to explain behavior that is both influenced by the current expected rewards and by past behavior.

To test whether a repetition bias is required at all to explain the behavioral data, we considered three alternative models that do not include an explicit repetition term. First, we used the expected value model (EVM), which posits that participants know the exact expected rewards and performed actions to solely maximize the expected rewards. However, as explained in the Methods section, this model would require infeasible computations made by participants as they perform the task.

Second, we used a model that is based on expected reward structure only. For this expected value with proxy model (EVPM), the reward is known for those sequences that have been chosen before, but for all others an approximated value *R*_0_ is used, which we assume participants estimated based on task instructions and training. Third, we considered the possibility that participants prefer the DAS based on the initial training and instructions. To model this, we used an extension of the EVPM, the expected value with proxy and default bias model (EVPBM), which has a constant bias in favor of the DAS to account for the observed high probability of DAS choices in our data. For details on the models, see Materials and Methods.

### Model Comparison

We calculated the predictive accuracy of the four cognitive models (EVPRM, EVM, EVPM, EVPBM) at the group level. We used the leave-one-out information criterion (LOOIC) (***Vehtari et al., 2017***) that evaluates model fit but also penalizes for model complexity (see Materials and Methods for details). Lower LOOIC values indicate a higher predictive accuracy, i.e., a lower difference between model predictions and observed data. We found that the EVPRM, the model including learning of repetition biases, showed the highest predictive accuracy (*LOOIC* = 69, 942.82, *SE* = 825.94) compared to the EVPBM (*LOOIC* = 73, 285.86, *SE* = 983.73), the EVPM (*LOOIC* = 74, 301.96, *SE* = 1, 018.95), and the EVM (*LOOIC* = 162, 308.58, *SE* = 1, 355.02) (see ***Table 2***. Because of its low predictive accuracy we excluded the EVM from further analyses.

**Table 2.** Results of model comparison

| Model | *LOOIC* | *SE* | *pLOOIC* | *dLOOIC* | *dSE* | Best Fit |
| --- | --- | --- | --- | --- | --- | --- |
| EVPRM | 69,942.82 | 825.94 | 810.92 | 0.00 | 0.00 | 27 |
| EVPBM | 73,285.86 | 983.73 | 1,054.58 | 3,343.05 | 382.35 | 15 |
| EVPM | 74,301.96 | 1,018.95 | 772.52 | 4,359.14 | 421.93 | 28 |
| EVM | 162,308.58 | 1,355.02 | 371.92 | 92,365.76 | 1,319.67 | 0 |

EVPRM: expected value with proxy and repetition bias model; EVPBM: Expected value with proxy and default bias model; EVPM: expected value with proxy model; EVM: expected value model; *LOOIC*: leave-one-out information criterion (lower values indicate higher predictive accuracy); *SE*: standard error of *LOOIC*; *pLOOIC*: effective number of parameters penalty; *dLOOIC*: *LOOIC* difference relative to the model with highest predictive accuracy, i.e. lowest *LOOIC* value; *dSE*: standard error of *dLOOIC* based on point-wise estimates; Best Fit: number of participants whose behavior was best explained by the model.

Following the guidelines from ***McElreath (2020***) for interpreting LOOIC values, we found that EVPRM described the data significantly better than the second-best model EVPBM: the standard error of the LOOIC differences *dSE* between EVPRM and EVPBM was substantially smaller than the difference in LOOIC between these models *dLOOIC* (see ***Table 2***). To investigate how well the models explained behavior at the participant level, we compared the LOOICs of the three remaining candidate models for each participant individually.

First, we counted how many participants were fitted best by each of the three candidate models.

This classification showed no clear pattern, as a considerable number of participants were equally well explained by each of the models (see ***Table 2***). Although EVPRM was the best model on the group level, the behavior of only 27 out of 70 participants (ca. 39%) was described best by EVPRM.

As a next step, we looked at the individual LOOIC values of the three models (see ***Figure 4***). Here, most of the participants whose behavior was described best by EVPRM showed a difference between the LOOICs of the candidate models, indicating that the EVPRM explained behavior better than the alternative models. In contrast, the LOOICs of those participants whose behavior was best described by the two alternative models did not show a clear difference in LOOICs. This indicates that the three candidate models explained behavior equally well. As the participants fitted best by EVPBM and EVPM had low repetition biases (see next section and ***Figure 5***), the EVPRM and the two alternative models are practically mathematically equivalent. We conclude that the EVPRM is the best model for 27 of the participants and is as good as the other two models for the remaining 43 participants.

**Figure 4.**
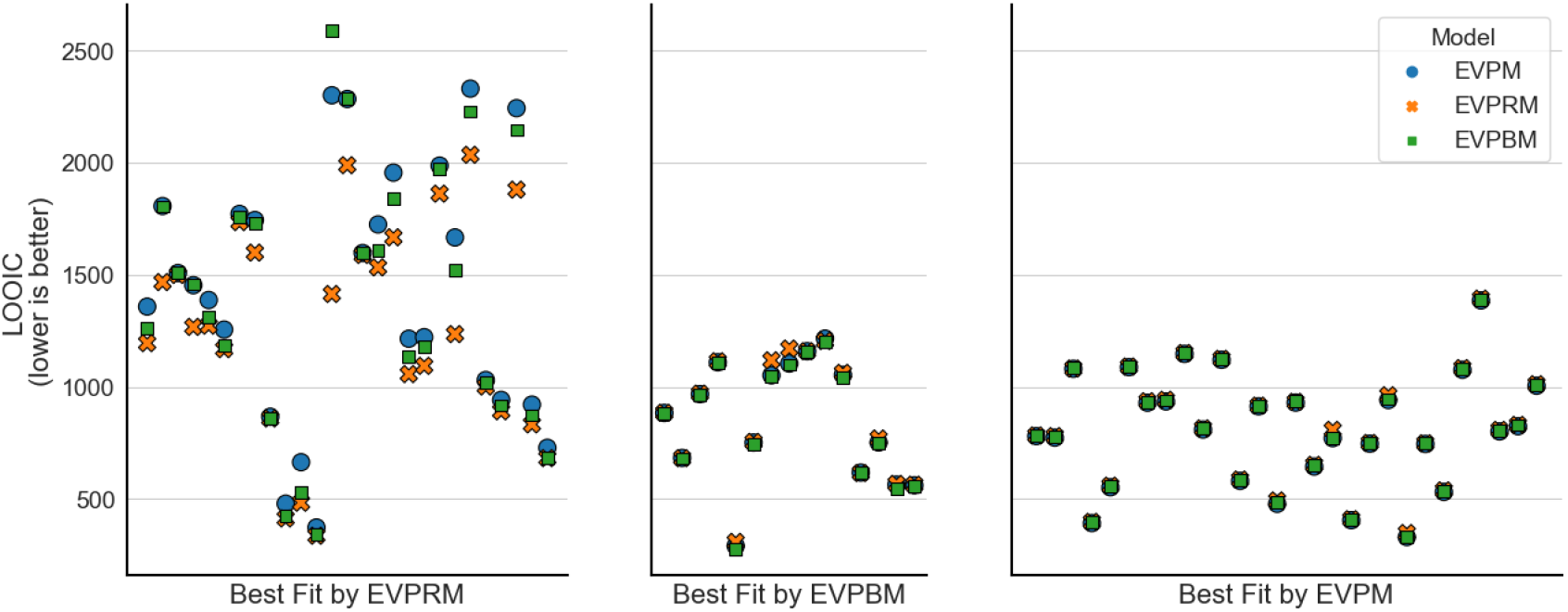
Model comparison at participant level. Predictive accuracy indicated by *LOOIC* for each participant and each model, with the three models for each participant aligned vertically. Participants are grouped depending on which model showed the highest predictive accuracy.

**Figure 5.**
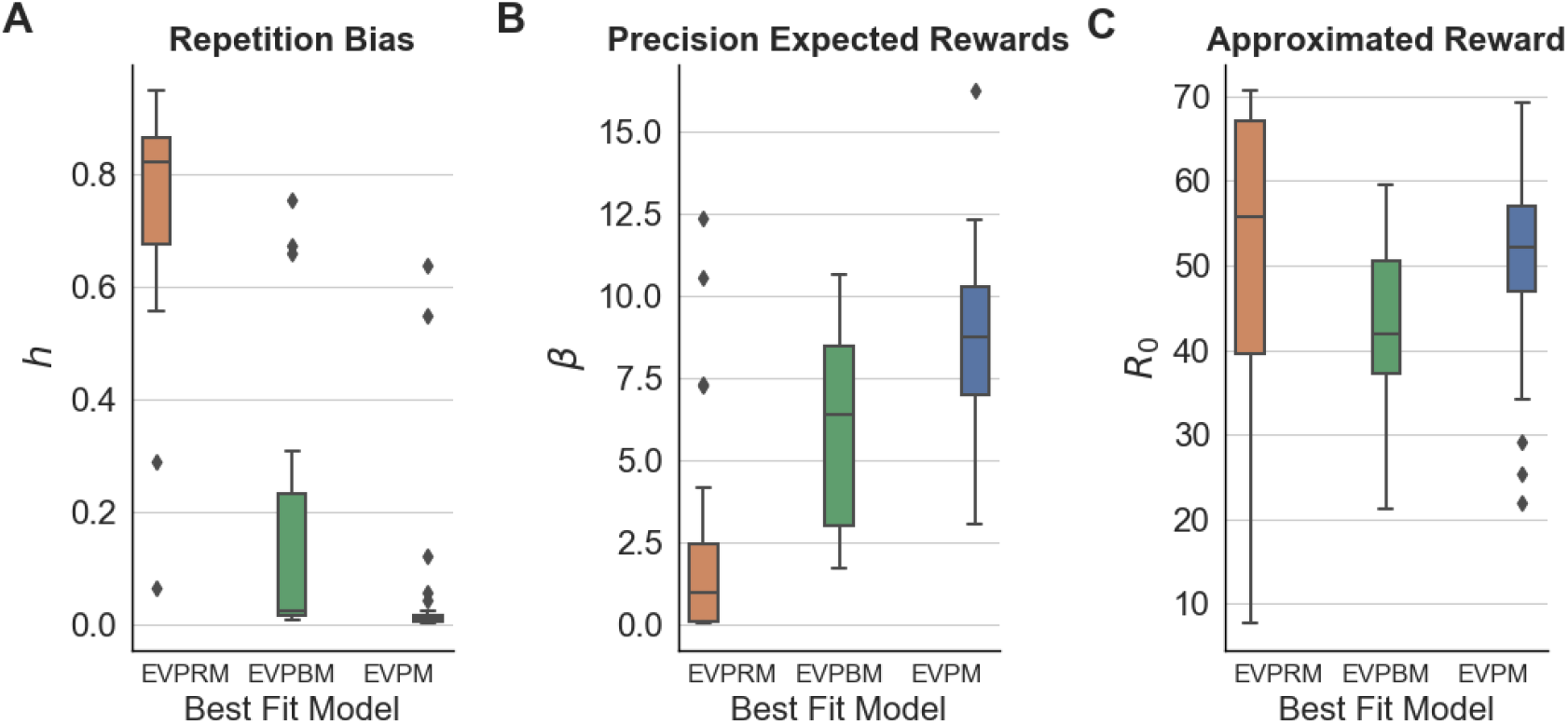
Estimated parameter values of EVPRM partitioned by the model that explained participant behavior best. Estimated value distribution for the three parameters of the EVPRM: **(A)** repetition bias *h*, **(B)** precision on expected rewards *fJ*, and **(C)** approximated reward *R*_0_. Participants are partitioned according to the model with the lowest *LOOIC*. Boxes represent the interquartile range (IQR). Horizontal lines inside boxes represent medians. Whiskers represent the 1.5 IQR of the lower and upper quartile.

Next we assessed if participants best fitted by EVPRM are the participants with a strong repetition bias. To do this, we analyzed the distribution of the inferred parameter values of the EVPRM and grouped participants based on the model that explained their behavior best (see ***Figure 5***). As expected, participants whose behavior was best explained by EVPRM showed the highest inferred repetition bias strengths. Furthermore, participants whose behavior was best explained by EVPM showed the highest inferred precision on expected rewards. The inferred parameter values of the approximated reward did not differ between best model fits.

In what follows, we compare fitted model parameters with behavioral measures of performance. Given that the EVPRM model has the best fit for 27 participants, and fits the remaining participants as well as the other models, we limit our analyses to EVPRM fitted parameters.

### Increase in DAS usage in participants fitted best with EVPRM

In our standard analyses above, we did not find a significant increase of DAS usage from the first to the second half of the experiment over all participants. We repeated this analysis with only those 27 participants best fitted by the EVPRM. This group of participants showed a high repetition bias (see ***Figure 5***), and as a consequence, we expected a significant increase of using the DAS from the first to the second half of the experiment for these participants. Indeed, the proportion of DAS choices of these participants significantly increased from the first half (46.2%, *SD* = 24.9%) to the second half (51.6%, *SD* = 26.1%) of the experiment, *t*(26) = −2.27, *p* = .02, *d* = .21.

### Correlations between parameters of EVPRM

We analyzed the correlations between the parameter estimates between the three free model parameters repetition bias *h*, precision on expected rewards *β*, and approximated reward *R*_0_.

As our model represents the influence of the repetition bias and expected rewards separately we can investigate the correlation between these two parameters. We expected that participants with a strong repetition bias *h* are potentially more guided by past behavior than by expected rewards. Therefore, precision over expected rewards and repetition bias strength should show a negative correlation. We found such a significant negative correlation between the precision over expected rewards *β* and the repetition bias strength *h, r* = −.75, *p <* .001 (see ***Figure 6***A). In addition, we found a significant positive correlation between *β* and the approximated reward *R*_0_, *r* = .30, *p* = .01 (see ***Figure 6***C).

**Figure 6.**
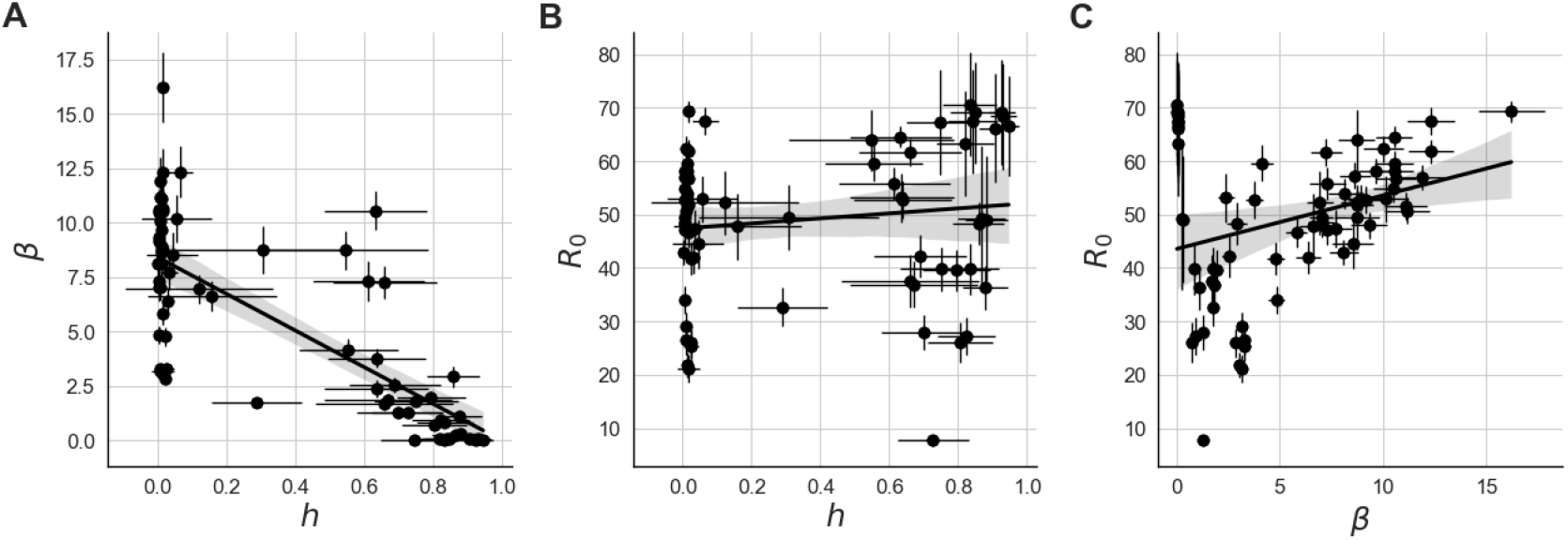
Participant-level correlations between estimated parameters of the EVPRM. **(A)** Correlation between repetition bias strength *h* and precision over expected rewards *fJ*. **(B)** Correlation between repetition bias strength *h* and approximated reward *R*_0_. **(C)** Correlation between precision over expected rewards *fJ* and approximated reward *R*_0_. Thick solid lines represent linear regression model fitted to the data. Thin solid lines represent standard deviations of individual fitted parameter values (*SD*).

One reason for a strong repetition bias might be a low approximated reward. Therefore, the expectation of a low reward for unobserved sequences of actions could lead to stronger action repetition if participants can find an alternative sequence with higher reward. Contrary to our prediction, the approximated reward showed a positive correlation with the repetition bias strength, but this correlation was not significant, *r* = .12, *p* = .30 (see ***Figure 6***B).

### Correlations between model parameters of EVPRM with performance measures

As a further intuitive validation measure, we also tested for correlations between the model parameters and performance measures. We expected that participants with a higher approximated reward *R*_0_ would show a decreased reliance on the DAS, due to an expectation of higher rewards for alternative sequences of action. These participants should deviate from the DAS more frequently. Inferred values of *R*_0_ correlated indeed negatively with the proportion of DAS choices *p*(DAS), but this correlation was not significant, *r* = −.18, *p* = .14 (see ***Figure 7***A).

**Figure 7.**
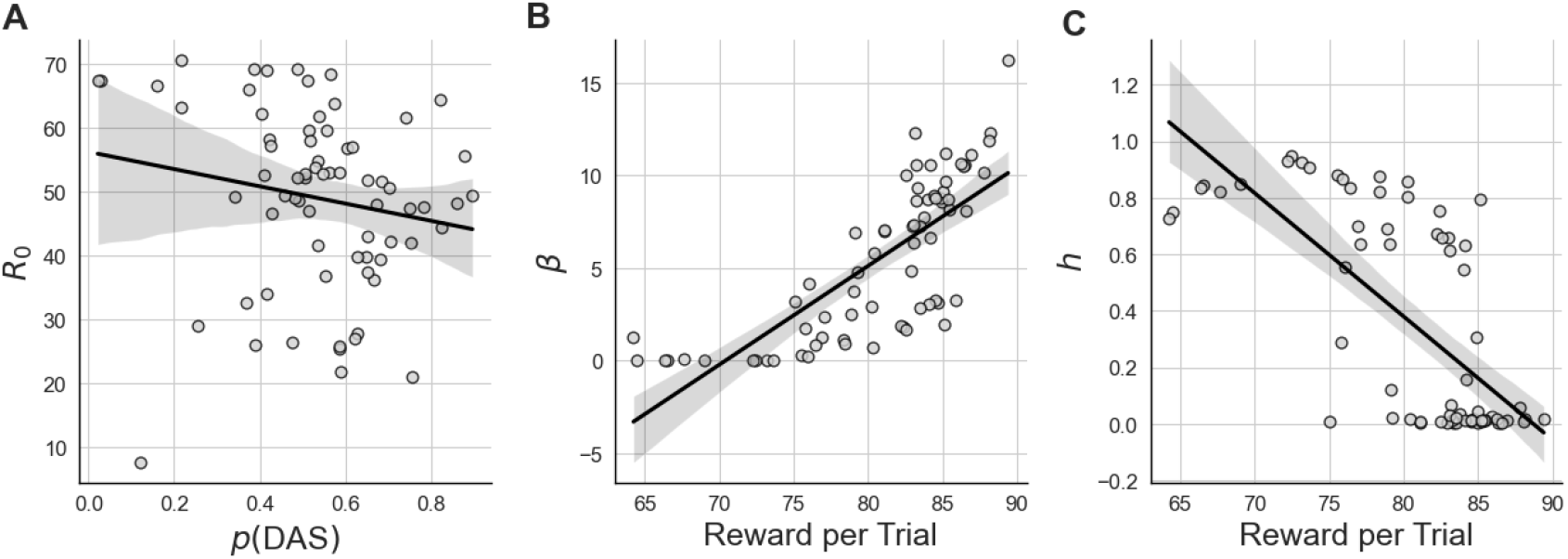
Correlations between estimated parameters of expected value with proxy and repetition bias model (EVPRM) and performance measures. **(A)** Correlation between free parameter of approximated reward *R*_0_ and the mean proportion of DAS choices *p*(DAS). **(B)** Correlation between model parameter of precision over expected rewards *fJ* and mean reward per trial. **(C)** Correlation between model parameter repetition bias strength *h* and mean reward per trial. Black solid lines represent linear regression model fitted to the data.

**Figure 8.**
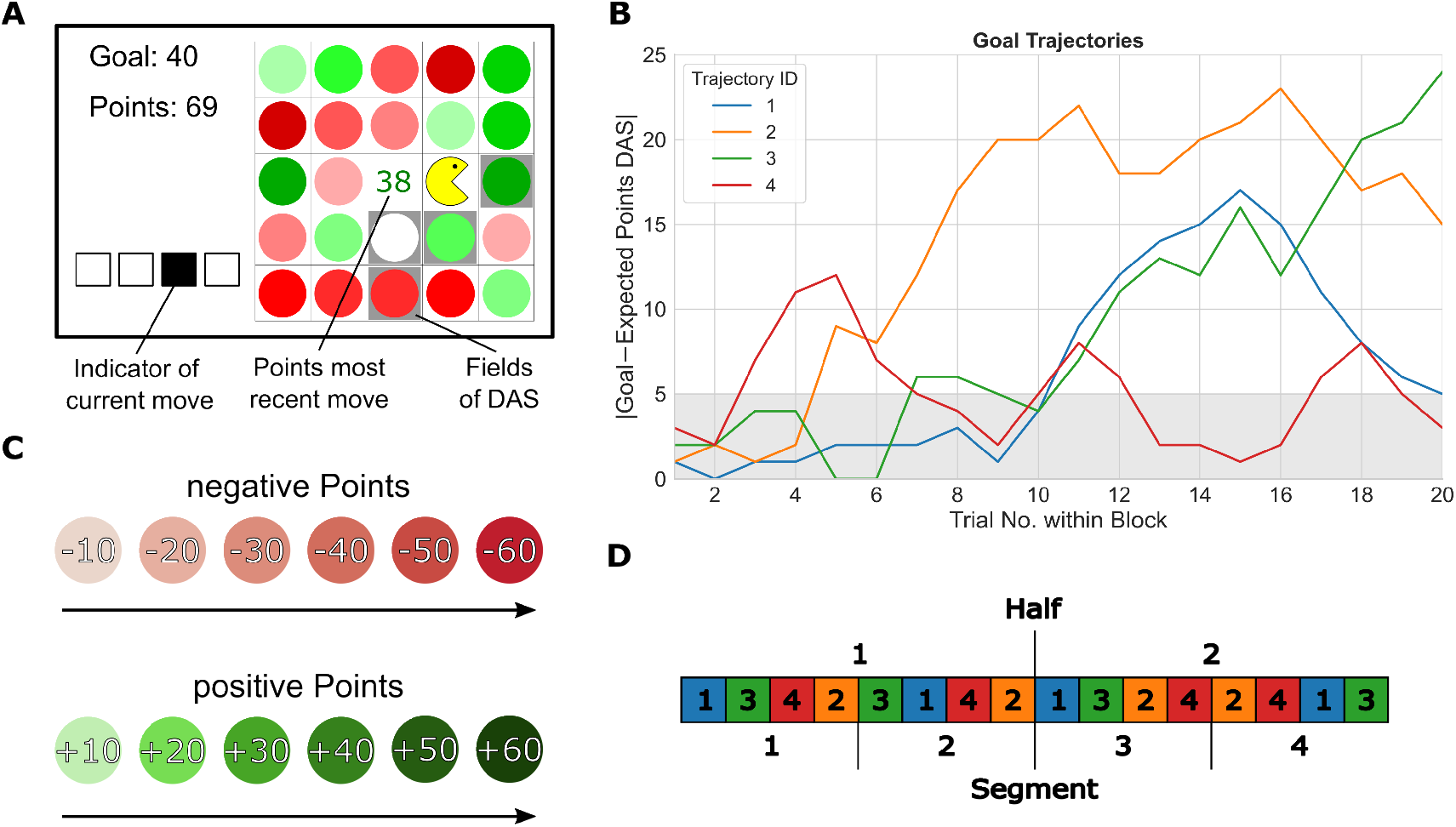
Experimental task. **(A)** Participants had to collect four colored circles by navigating a Pacman-style character on a 5-by-5 grid. Each circle’s color indicated the number of points obtained when moving into the corresponding field (see main text for details). The points obtained for the most recent collected circle were displayed in the centre of the grid (here 38). The goal of the current trial was presented in the top-left corner. Below this goal number, the current sum of points gathered up to the current move were displayed. Four squares located at the bottom left indicated the current move number. In the example, the participant was about to select its third move after having collected points from two circles. The default action sequence (DAS) fields were highlighted with a gray background. **(B)** Graph of the four goal point trajectories used. Each block consisted of 20 trials. The y-axis shows the absolute difference in points between the goal point per trial, and the expected points one would obtain by using the DAS. The gray area represents trials where the DAS was one of the available action sequences with the highest expected reward. **(C)** Visual representation of points per color. **(D)** Order of goal trajectories within the experiment and mapping of blocks into halves and segments.

**Figure 9.**
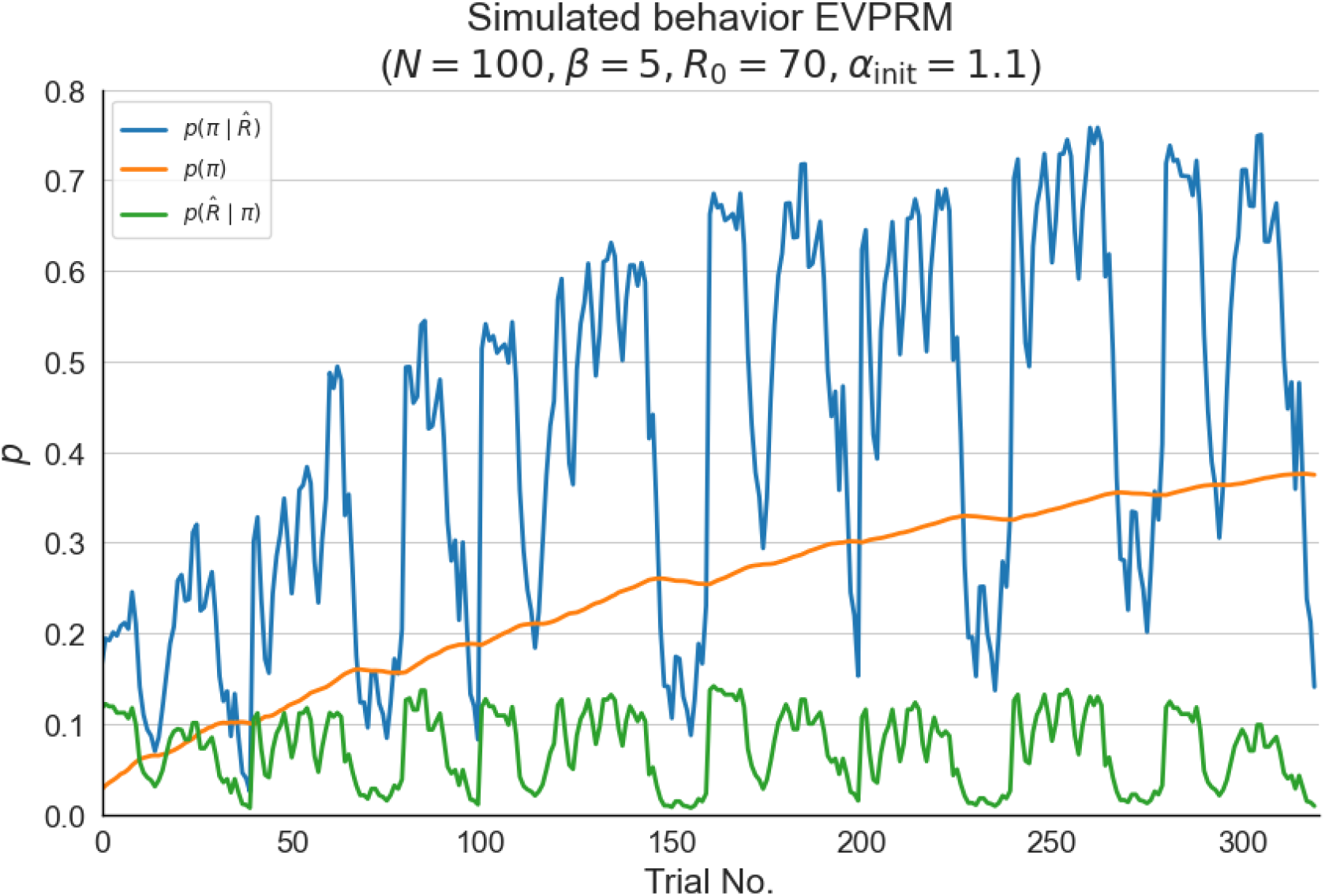
Simulation of task with EVPRM. Means for probability of selecting one specific sequence of actions *π*, 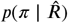 (blue line), the prior over policies for *π, p*(*π*) (orange line), and the expected reward for *π*, 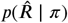 (green line) over *N* = 100 simulations. While expected rewards are in a constant range throughout the task, the prior over policies increases and accordingly the choice probability.

Further, we expected that participants with higher precision over expected rewards *β* were likely to earn more reward. This is because as *β* increases, participants would have a lower uncertainty on the expected rewards. This increases the probability that participants select actions with higher expected rewards. As expected, participants showed a positive significant correlation between *β* and the mean reward per trial, *r* = .76, *p <* .001 (see ***Figure 7***B).

Crucially, we expected that participants with higher repetition bias strength *h* would receive lower rewards because participants with a strong repetition bias tend to repeat past behavior rather than to maximize expected rewards. We found this significant negative correlation between the repetition bias strength and the mean reward obtained per trial, *r* = −.69, *p <* .001 (see ***Figure 7***C). Accordingly, the achieved reward decreased with increasing repetition bias strength.

We also expected that participants with stronger repetition bias *h* show shorter reaction times (RTs), as the repetition of past behavior should be executed faster than selecting yet unknown sequences of actions. Contrary to our expectation *h* showed a significant positive correlation with mean RTs, *r* = .37, *p* = .001. We speculate that participants with a strong repetition bias were probably not as motivated as other participants and therefore slower in processing relevant stimuli and/or executing the movements. In combination with the tight deadline of six seconds, these participants probably relied more strongly on known sequences of actions. This speculation is supported by the significant positive correlation between the number of time out trials and the repetition bias. See Appendix ***Table 2*** for all correlational analyses.

## Discussion

In this study, we have shown experimental evidence of a repetition bias that increases the probability of performing a sequence of actions as a function of how frequent this action sequence had been used before, over the course of the experiment. To show this repetition bias, we introduced a new grid-world task and employed a recently proposed computational model that describes action selection as a balance between goal-directed and a repetition bias. We found that the repetition bias was negatively related to task performance, suggesting an opposition between goal-directed performance and repetition bias.

We developed the Y-navigation (Y-NAT) task where participants had to meet trial-specific goals by collecting points. We gave participants information about a default action sequence (DAS) that let them obtain the maximum expected reward in nearly half of all trials. With this manipulation we ensured that participants repeat at least one sequence of actions frequently.

In our behavioral analyses, we found that nearly all of our participants used the DAS most frequently. Interestingly, participants executed the DAS even when the DAS did not provide the highest reward. However, our subsequent standard analyses to test for a repetition-induced increase in DAS usage only revealed non-significant trends, and the evidence remained inconclusive.

We complemented our analyses by a computational modeling approach to assess whether explicit modeling of repetition learning over the course of the experiment reveals a repetition bias. Indeed, the repetition bias model explained participants’ behavior best.

With the proposed model-based approach we were able to rule out several alternative explanations for the observed effects. First, we can exclude random responding, as all participants either relied on expected rewards or showed a repetition bias. Second, we excluded behavioral repetition as a fixed-choice strategy not influenced by past actions. For instance, one strategy could be to always select the incentivized DAS. As the DAS was of high value during the initial few trials of each block and always resulted in a reward, consistently repeating the DAS would classify as goal-directed behavior and would not be evidence for a repetition bias. We tested this alternative using a model that replaced the repetition bias effect by a constant bias added to the expected reward for the DAS, leading to constant higher choice probabilities for the DAS over the course of the experiment. Using model comparison we ruled out this alternative. Further evidence against a constant but not increasing influence of repetition is the finding of a general reduction in behavioral variability over time.

The finding that the behavior of only a subgroup of participants was best explained by the repetition bias model seems consistent with previous studies where only subsets of participants were found to show habitual behaviour (***Pool et al., 2022; Gera et al., 2023***). One reason might be a strong motivation to perform well in experimental tasks (***Cerasoli et al., 2014***). This motivation probably prompts participants to use goal-directed behavior to collect reward. This effect might be strengthened by our performance-based bonus payment, as incentives have been shown to modulate cognitive effort (***Patzelt et al., 2019***). This interpretation is also consistent with the finding that, contrary to our prior expectation, participants with an increased repetition bias showed slower reaction times. Slower reaction times are related to poorer performance and could be an indicator of less motivation and thus less goal-directed behavior for participants with strong repetition bias.

### Repetition bias and cost-benefit arbitration

In our task, the effect of the repetition bias can only be measured in combination with concurrent goal-directed behavior. Specifically, according to the model, the first few decisions in a new task context are mainly based on expected rewards. Concurrently, the effect of the repetition bias ramps up and has, as we found, a measurable effect on action selection. While this is the concrete mechanism in the present model (see also ***Schwöbel et al. (2021***)) the increasing influence of the repetition bias could also be viewed as an efficient, dynamic cost-benefit arbitration.

Model-based planning is associated with cognitive costs (***Shenhav et al., 2013; Kool et al., 2017***), and it has been postulated that decision makers compute whether it is worth investing the cognitive effort. It might be that the inferred repetition bias strength is just a measurable expression of such a cost-benefit arbitration.

An alternative view is to turn this argument around and to postulate that the computation and use of the repetition bias is the causal underlying mechanism, which is observed and eventually interpreted as an apparent dynamic cost-benefit analysis. What speaks for this view is that the repetition bias is simple to compute because the model just increases a task-specific counter by 1. In the brain, this would correspond to a simple strengthening of a context-action association. Conversely, it has been shown that, in principle, cost-benefit arbitration leads to a computationally involved recursive planning process (***Shenhav et al., 2013***). The question, which can be addressed in future studies, is whether a simple repetition bias computation is enough to explain apparent cost-benefit computations to generate behaviour.

### Relation to other models and implications for habit learning

The idea of a repetition bias is well established in psychology (***Thorndike, 1911***). It is related to stimulus-response (S-R) learning, as repetition facilitates the formation of S-R associations and re-cency effects in value-based decision-making tasks (***Guthrie, 1952; Wood and Rünger, 2016; Watson et al., 2022***). The repetition bias is also consistent with previous proposals for the role of action repetition in the development of habitual behavior (***Thorndike, 1911; Miller et al., 2019; Schwöbel et al., 2021; Nebe et al., 2024***). Similarly, behavioral repetition of action sequences has been identified as a way to optimize the trade-off between maximizing reward and a reduction of policy complexity (***Gershman, 2020***).

Importantly, the repetition learning mechanism is different from stimulus-response associations typically found in devaluation studies (e.g. ***Horstmann et al., 2015; Dickinson et al., 1983; Hardwick et al., 2019***), and different from a potential trade-off between model-free and model-based reinforcement learning (RL) (***Daw et al., 2011; Dolan and Dayan, 2013***), because, in contrast to model-free RL, the repetition bias is value-free (***Miller et al., 2019***) and does not directly depend on past rewards.

The repetition bias is possibly a prerequisite for the development of habitual behavior (***Wood and Rünger, 2016; Miller et al., 2019; Schwöbel et al., 2021; Nebe et al., 2024***). This opens up possibilities to use this mechanism and its predictions to investigate the formation of habits. Especially, concerning the lack of a unified methodology for measuring habits (***Watson and De Wit, 2018; Watson et al., 2022***), our task and the repetition bias could in principle be used to measure the tendency towards habitual behavior during ‘only’ a few hundred trials without the need to implement habitual learning with over thousands of trials (***Hardwick et al., 2019; Luque et al., 2020; Frölich et al., 2023***) and sessions over two (***Frölich et al., 2023***) to four (***Hardwick et al., 2019***) days.

Indeed, many studies investigated the influence of repetition through habits. In these studies habits are typically only measured indirectly, as a lack of goal-directed behavior during an extinction phase (***Balleine and Dezfouli, 2019; Watson et al., 2022***). However, a lack of goal-directed behavior can alternatively emerge due to an inaccurate representation of action-outcome contingencies during extinction, or random responding due to a lack of motivation (***Watson et al., 2022***). Instead, here we measured repetitive behavior directly through a combination of task design and a model-based approach, enabling us to measure positive characteristics of repetitive behavior. Additionally, our task did not consist of separate training and extinction phases, and we provided feedback after each trial. This approach avoids a potentially inaccurate representation of the expected rewards.

### Conclusion

In conclusion, we introduced a novel sequential decision making task, where we demonstrated the influence of both expected rewards and a repetition bias on decision making. Using computational modeling we provided empirical evidence for a repetition bias which is simply expressed as a value-free increase of choice probability each time an action is performed. This repetition bias mechanism may underlie habit formation and emphasizes the importance of considering frequency-based mechanisms besides reward-driven mechanisms in future studies.

## Materials and Methods

### Participants

Participants were recruited using the recruitment system of the faculty of psychology at the TUD Dresden University of Technology. In this system, students and individuals from the general population interested in being participants in psychological studies can register. 74 participants completed the experiment. Four participant were excluded for lack of behavioral variability (they performed the same sequence of actions in more than 90% of all trials). The remaining 70 participants (50 female) had a mean age of 24.1 years (*SD* = 4.6). All participants confirmed that they did not have dyschromatopsia.

Remuneration was a fixed amount of 10€ or class credit plus a performance-based bonus (*M* = 2.58€, *SD* = 0.19€). The bonus was determined as a linear function from each participant’s rewards acquired during the experiment, where a reward of 100 yielded 1ct. Participants were informed about the maximum of the bonus, but not the exact calculation.

The study was approved by the Institutional Review Board of the TUD Dresden University of Technology (protocol number EK 578122019) and conducted in accordance to ethical standards of the Declaration of Helsinki. All participants were informed about the purpose and the procedure of the study and gave written informed consent prior to the experiment.

### Experimental task

Data collection was performed online. The task was built using lab.js (***Henninger et al., 2021***) and hosted on the neurotests server of the TUD Dresden University of Technology, which is specifically designed for hosting lab.js tasks.

Participants had to navigate a Pacman-like character across a 5-by-5 grid using their keyboard (see ***Figure 8***A) to collect points matching a pre-defined trial-specific goal. In every trial, participants had to execute a sequence of four actions within a time limit of 6s. The action-set was restricted to moves in three directions: diagonally to the upper left, diagonally to the upper right or directly downwards. This specific choice of navigation, inspired by the work of ***Fermin et al. (2010***), was designed to restrict the available sequences of actions participants could take. Exiting the grid’s boundaries or revisiting a previously visited field was not possible. Any attempt to do so triggered a red warning message, requiring the participant to redo the move.

Upon each action, the character visually moved to the designated field and thereby collected the circle within that cell. Circles were colored to represent point values: Green circles represented positive points ranging from 10 to 60 in increments of 10, while red circles represented negative points ranging from −10 to −60. The shading of the color indicated the magnitude of points, with darker shades representing higher positive or negative values (see ***Figure 8***C). Additionally, a Gaussian distributed noise (with *µ* = 0, and *(5* = 1.3) was applied to the points earned from each move and the resulting value was rounded to the nearest integer. After each move, the points from the collected circle were displayed at the center of the grid, and the sum of points collected during that particular trial was displayed at the top left corner (see ***Figure 1***). The trial’s total score was calculated as the sum of points from the combined sequence of four actions.

Importantly, the main goal of the task was to match the trial’s points—achieved from the sequence of four actions—as closely as possible with a predefined, trial-specific goal. Participants’ reward for each trial was then calculated based on the difference between the trial’s total points and the predefined goal. Smaller differences (in absolute value) led to higher rewards:

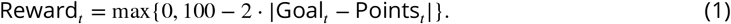

The received reward was displayed as a green bar at the end of each trial in the feedback stage (see ***Figure 1***). If participants did not complete four moves within the time limit, no reward was earned and a red warning message appeared at the feedback stage on the left side. The sum of collected rewards defined the performance-based bonus payment.

Importantly, to ensure that participants repeat at least one sequence of actions frequently, we introduced a default action sequence (DAS). This sequence was visually indicated by a grey back-ground color of the four corresponding fields (see ***Figure 8***A) and participants were given information about the average number of points that could be collected with the DAS at the start of each trial at the center of the grid (see the left-most panel of ***Figure 1***). The fields, sequence of actions, and sum of collected points of the DAS were the same throughout the experiment. In the example of ***Figure 1***, the DAS comprised two downward moves first, followed by two consecutive up-right movements.

The experiment was divided into 16 blocks consisting of 20 trials each. The distribution of points remained constant within a block, but changed between blocks. The points of the four circles of the DAS also differed across blocks but the sum of the points remained constant. The block sequence of the distributions of points were the same for each participant.

Within a block, goal points changed over trials. We used four different trajectories of 20 goals each (see ***Figure 8***B). Goal trajectories differed by their maximum difference between the goal points and the expected points of the DAS, and their trend of this difference throughout the block. This difference ranged between 12 and 24 points. Note, that while the points that could be collected with the DAS stayed the same, the goals changed between trials, and therefore the rewards for the DAS varied.

We selected these four trajectories to represent different principled trajectories that would it make diffcult for participants to predict whether the DAS would remain optimal during the duration of a block. All four trajectories started out close to the expected points obtainable by the DAS. Only later into the block goal points started to deviate from this initial value, or not. For example, one goal trajectory was remaining close to points obtainable by the DAS while another one increased only after half the block but then decreased again.

With this procedure we effectively proposed an optimal sequence of actions at the first few trials of each block that was slowly devalued during subsequent trials. A repetition bias should manifest by increased DAS choices over the course of the experiment for the same expected rewards and should be detectable with summary statistics.

We subdivided the blocks into four segments of four blocks each (see ***Figure 8***D). Within each segment all four trajectories of goals were used once in a pseudo-random order so that no trajectory was repeated in two consecutive blocks. This order of goal trajectories was the same for each participant.

To promote the use of the DAS, we manipulated three features of the task. First, at least the first two trials of each block had a trial-specific goal close to the points of the DAS (see ***Figure 8***B). Overall, in 43.75% of all trials, the absolute difference between the trial goals and the expected points of the DAS was between zero and five points. Therefore, for these trials, due to the minimum difference of ten points between circles of different color, the DAS was one of the available action sequences with the highest expected reward. In the remaining trials, there was always at least one sequence with a higher expected reward than the DAS.

Second, we only used a partial devaluation of the DAS: the lowest expected reward of the DAS was still about half of the maximum reward. This follows from the reward calculation (see ***Equation 1***) and the maximum difference between goals and the expected DAS points of 24 (see ***Figure 8***B).

Third, using the DAS gave participants a probabilistic bonus reward of 20 during half of the blocks. These bonus blocks were distributed pseudo-randomly throughout the experiment to ensure that the bonus was available always during two of the four presentations of each goal point sequence. This probabilistic bonus could be earned for each trial within a bonus block, but exclusively when using the DAS. The probability of receiving the bonus was *p* = .25, although the precise probability remained undisclosed to the participants. Participants were informed about upcoming bonus blocks right before they started. In addition, during the trial start phase, bonus trials were indicated by changing the color of DAS points from gray to blue. A blue bar next to the green reward bar during the feedback stage indicated the receipt of the bonus.

The experiment started with an elaborate training phase to ensure that participants understood the task. The first part involved an introduction to the navigation, which was followed by familiarizing participants with the color coding of the circles. Then trial-specific goals and reward calculation was introduced. This was followed by introducing the DAS, and finally the bonus factor was explained. During this part of the training participants had to meet no deadline and could spend as much time as they needed.

After this introductory phase, participants practiced two blocks as they would appear later in the main experiment. One block was with probabilistic bonus and one without. The only difference to the main experiment was an extended deadline of 10s.

Between blocks, participants had the opportunity to take a self-determined break. The experiment, including training, had a total duration of approximately 60 minutes. The performance-dependent bonus rewards were determined by adding the rewards of all trials in the main experiment.

Data analysis was performed in Python using the packages NumPy, Pandas, ArviZ, Scipy’s Stats module, and pingouin. Task code, data and analysis code are publicly available at GitHub.

### Expected value with proxy and repetition bias model (EVPRM)

We made use of a previously published repetition-based learning model, the prior-based control model (***Schwöbel et al., 2021***), for model-based data analyses. This model describes a mechanism for taking into account previous action sequences when making choices. The model counts how many times each action sequence *π* has been chosen in the past. This contributes to the decision-making process as a prior distribution over policies *p*(*π*) that represents the probability of selecting an action regardless of expected reward or any other task contingency: the influence of the prior over policies on action selection of a specific action increases depending on how many times this action has been chosen before. The model is complemented by a component 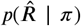 based on expected rewards given the predicted outcomes of the performed actions (i.e. value-based). These two components play the role of priors and likelihood, respectively, to turn decision-making into a Bayesian inference process:

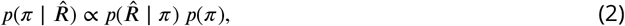

where 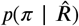 represents the posterior distribution that is defined by the probability of choosing policy *π* given the reward structure 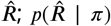 represents the expected reward 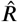 given policy *π*; and *p*(*π*) is the prior over action sequences. The multiplication of the expected reward and the prior over policies balances the influences of goal-directed planning and past behavior on action selection. Intuitively, the goal-directed component 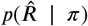 represents the value-based part of making a decision, i.e. a participants simply selects the action that gives the highest expected reward, while the prior over action sequences *p*(*π*) implements the repetition bias.

In our experimental task, the reward structure 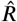, i.e. the expected reward for each action sequence *π*, can in principle be calculated given the information available to participants: the points for each one of the squares on the grid is shown on the screen, so participants could calculate the points of every of the possible 36 sequence of actions *π* and determine the expected reward with the exception of a noise term that is not influenced by *π*. However, as there is a deadline of six seconds, the calculation becomes unfeasible. To account for this, we posit that participants rely on prior beliefs or approximations they might have acquired in previous trials.

For the proposed EVPRM, we assumed a reward structure 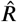 that depends on past observations made by the participant: for sequences that have been already observed during the current block, the model uses the exact observed reward; for the unobserved sequences, it uses an approximated reward *R*_0_, which we assumed participants approximate based on their experiences during previous blocks and training. The approximated reward *R*_0_ is a free parameter and indicates the individual expected reward for all yet unobserved sequences of actions. As an exception, the DAS was always assumed to be an observed sequence of actions because the points of the DAS were communicated at the initial phase of each trial. Also the expected reward of the DAS included the probabilistic bonus reward. With this, the reward structure in the EVPRM is as follows:

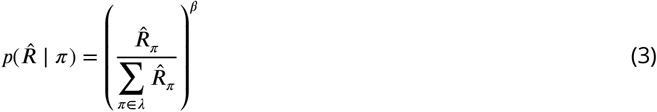

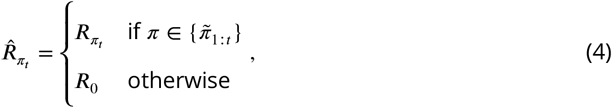

where *π* = {(*a*_1_, *a*_2_, *a*_3_, *a*_4_) I *a*_*i*_ ∈ {↖, *↓,↗* }}, with *π* = (*a*_1_, *a*_2_, *a*_3_, *a*_4_) represents a sequence of four actions, and *a*_*i*_ ∈{ ↖, *↓,↗*} represents the three movement directions, 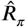 is the expected reward of the sequence of actions *π*,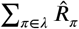 is the sum of expected rewards of all sequences of actions, with *-λ* representing all possible action sequences, *β* is a free parameter representing the precision over expected rewards, *R*_*π*_ is the expected reward for the sequence of actions *n, R*_0_ is the approximated reward for unobserved sequences of actions, 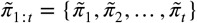 are the performed sequences of actions up to trial *t*. Note that we chose 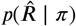 as a fraction to stay close to the Bayesian framework in ***Schwöbel et al. (2021***) and to have a comparable equation to the prior below.

The free parameter *β* represents the precision over expected rewards: values of *β >* 1 leads to more concentrated probabilities that favor the choice of the sequences of actions with the highest expected rewards and values of *β <* 1 lead to more uniformly distributed probabilities, enabling greater exploration of different choices.

The prior over action sequences *p*(*π*) was defined, as by ***Schwöbel et al. (2021***), by a counter *γ* for the number of times the respective sequence of actions has been used used in the past, and the initial count *α*_init_ is a free parameter that was equal for each sequence of actions:

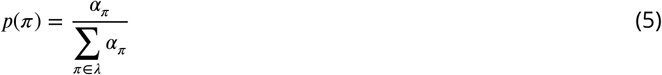

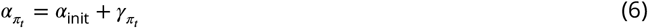

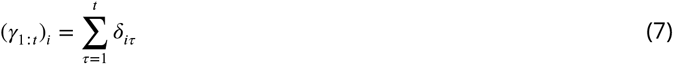

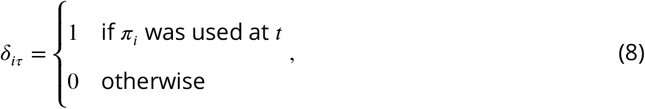

where 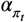 is the repetition bias strength at trial *t, α*_init_ is the initial count, 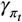 is the counter of how many times the sequence of actions *π* was performed until trial *t*, 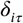 is the Kronecker delta.

Following ***Schwöbel et al. (2021***), the free parameter initial count *α*_init_ influences the strength of the repetition bias. A low initial count, e.g. *α*_init_ = 1, leads to a strong repetition bias. As *α*_init_ defines the counter for all sequences of actions, the increase of *γ* by 1, after a sequence of actions was performed, leads to a substantial increase of the prior over policies for this sequence. In contrast, a high initial count, e.g. *α*_init_ = 100, leads to a weak increase of the prior over policies if a sequence of actions is performed.

Finally, our model can make decisions at every trial by sampling from the categorical posterior probability distribution over possible *π*, defined as: 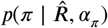, which is the probability of sampling each sequence of actions *π*, at each trial depending on the assumed reward structure 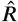, and the prior over policies *α*_*π*_ :

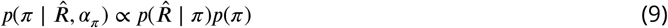

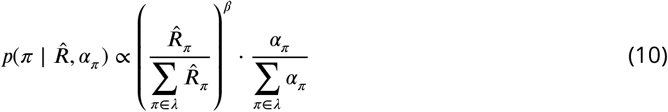

By changing the free parameters, we can change the behavior of the agent: At one end, with a high initial count *α*_init_, an agent will be minimally influenced by its past behavior and is nearly completely goal directed. At the other end, with a low initial count *α*_init_, agent behavior is more influenced by expected rewards and thus has a strong repetition bias of past action sequences. In addition, a precision over expected rewards *β* close to 0 represents the case in which the agent is uncertain about the learned reward structure and will tend to choose behavior based on the repetition bias.

In ***Figure 9*** we simulate an experimental session with our model, focusing on one action sequence *π*, the DAS. In the simulations, the model has a high influence of past behavior (*α*_*init*_ = 1.1). The used precision over expected rewards (*β* = 5) moderately pronounced the distribution of expected rewards. Based on the changing goals the expected reward 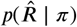 for this action sequence changes in a constant range from trial to trial throughout the experiment. But the prior over policies *p*(*π*) for this action sequence increases slowly over time, because this action sequence is performed repeatedly. One can see that in trials where the expected reward is relatively high, the resulting posterior 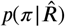 is high as well. This means the resulting choice probability is driven by expected rewards, represented by the first term on the right-hand side of ***Equation 10***. In addition, as the prior is slowly increasing, there is a growing contribution of the repetition bias, given by the second term of ***Equation 10***. Hence, the repetition bias increases the choice probability but actions are in principle still modulated by expected rewards. Effectively, in the example, the repetition bias increases the choice frequency from roughly 0.3 in the first 50 trials to roughly 0.7 in the last 50 trials, when there is a relatively large expected reward.

Note that the original model by ***Schwöbel et al. (2021***) was formulated within a planning as inference (***Botvinick and Toussaint, 2012***) and active inference (***Friston et al., 2016; Schwöbel et al., 2018***) framework to calculate the posterior distributions for action selection. We adapted the key idea: the posterior probabilities are based on the product of a function over the expected rewards and the prior over policies. Here, for our purposes, we simplified this model to derive a relatively straightforward observation model so that we could use Bayesian inference for fitting the model’s free parameters to participant data. Furthermore, the model calculates probabilities based on past and current observations and does not use any kind of future forward planning. It is therefore related to RL models, which also calculate subjective values for the possible actions based on the current expected rewards and the reward history (***Dezfouli and Balleine, 2012; Daw et al., 2011; Miller et al., 2019***).

### Alternative models

The proposed EVPRM makes two assumptions: (1) the repetition bias influences action selection, and (2) participants used an approximated reward for unobserved sequences of actions. To test against alternative explanations, we formulated three alternative models. These models differ only in their assumed reward structure 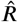 and as a critical distinction, they do not include the prior over policies *p*(*π*). In what follows, we introduce the three alternative models.

#### Expected value with proxy and default bias model (EVPBM)

An alternative explanation for the repetition of the DAS would be a bias specifically for the DAS but not a general repetition bias as formulated in the EVPRM. To implement this assumption, we derived a new model variant that had an exclusive and constant bias for the DAS. In other words, this model assumes that during training participants developed a bias for choosing the DAS but did not have a slowly increasing repetition bias or a preference for repeating other action sequences. Such a constant bias in favor of the DAS would also lead to an increase in DAS choices and be interpreted as a repetition bias. The difference to the EVPRM is that a constant bias would be independent from past behavior in the main experiment.

To model this bias, we added a constant term as a free model parameter to the expected reward of the DAS:

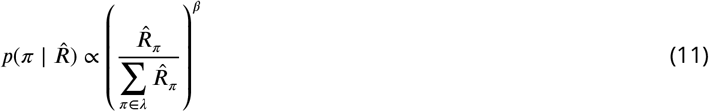

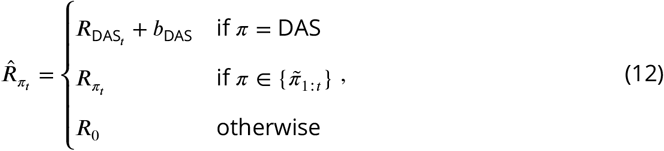

where *π* is a sequence of four actions defined as above, 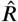 is the assumed reward structure,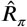 is the expected reward for the sequence of actions *π, β* is the free model parameter representing precision over expected rewards, *R*_0_ the free model parameter of approximated rewards for yet unobserved sequences of actions, 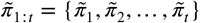 are the performed sequences of actions up to trial *t*, and *b*_DAS_ is the bias for *π* _DAS_.

#### Expected value with proxy model (EVPM)

A second alternative explanation of the choice data is that repetition does not influence action selection at all. Therefore, contrary to the EVPRM, participants’ behavior is not affected by past behavior, but determined by expected rewards only. To implement this assumption, we derived a model variant by removing the prior over policies from the EVPRM to have a model that is solely dependent on the expected reward structure:

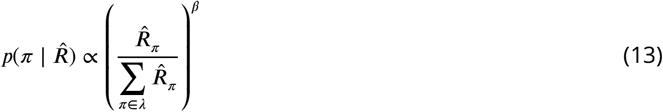

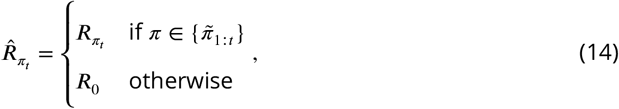

where *π* is a sequence of actions defined as before, 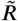 is the assumed reward structure, 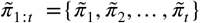 are the performed sequences of actions up to trial *t, β* is the precision over expected rewards and a free model parameter, *R*_*π*_ the expected reward for a sequence of actions *π*, and *R*_0_ the free model parameter of approximated reward for unobserved sequences.

#### Expected value model (EVM)

The EVPM relies on the approximated reward for unobserved sequences of actions *R*_0_. An alternative is that participants indeed were able to calculate expected rewards for all sequences of actions. To implement this assumption we instantiated a model without the approximated reward parameter and to use expected reward *R* _*π*_ instead. Therefore, contrary to the other candidate models, this model performs actions selection independent from past behavior. We implemented this as:

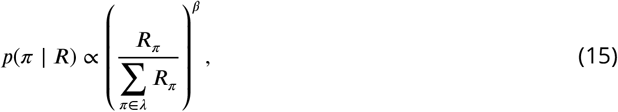

where *π* is a sequence of four actions defined as before, *R*_*π*_ is the expected reward of the sequence of actions *π*, and *β* is the free model parameter representing precision over expected rewards.

### Model fitting

Parameter estimation was done in Python with PyMC (***Salvatier et al., 2016***, version 5.0.1) using the No U-Turn Sampler (NUTS) (***Hoffman and Gelman, 2014***). We obtained 4,000 samples from four chains of length 1,000 (1,000 warm-up samples).

We used the following weakly informative prior distributions for the free model parameters: *β* ∼ Г(3, 1), *R*_0_ ∼ Г(55, .75), 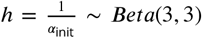, and *b*_DAS_ ∼ Г(3, .1). We used the same priors for all candidate models. The complete code can be found online at GitHub.

### Model comparison

To ensure that parameter inference works well for a meaningful range of parameters, we performed extensive parameter recovery studies for all four models (for details see Appendix ***Figure 1***).

Model comparison was based on using leave-one-out cross-validation approximated by Pareto-smoothed importance sampling (PSIS-LOO) (***Vehtari et al., 2017***). This information criterion calculates the pointwise out-of-sample predictive accuracy from a fitted Bayesian model. Crucially, it penalizes models with more parameters. We calculated the expected log point-wise predictive density (elpd) and the corresponding standard error (*SE*) on the deviance scale (−2elpd). Lower values of PSIS-LOO indicate higher predictive accuracy. Calculation of PSIS-LOO scores was performed with ArviZ (***Kumar et al., 2019***, version 0.7.0).

### Parameter distributions of EVPRM

To better compare individual repetition bias strengths we used the inverse of the initial count 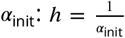 (see ***Equation 6***). *h* has a value range from 0 to 1, where values near 1 indicated a strong repetition bias and values around 0 indicate a weak repetition bias.

Repetition bias strength varied from very low values between close to 0 and .2 to medium to strong values between .5 and .9 (see ***Figure 10***A). The inferred *β* values (the precision over expected reward) spread between values very close to 0 and high values up to 16 (see ***Figure 10***B). The approximated reward *R*_0_ for unobserved sequences of actions showed a broad range of values between around 10 and around 70 (see ***Figure 10***).

**Figure 10.**
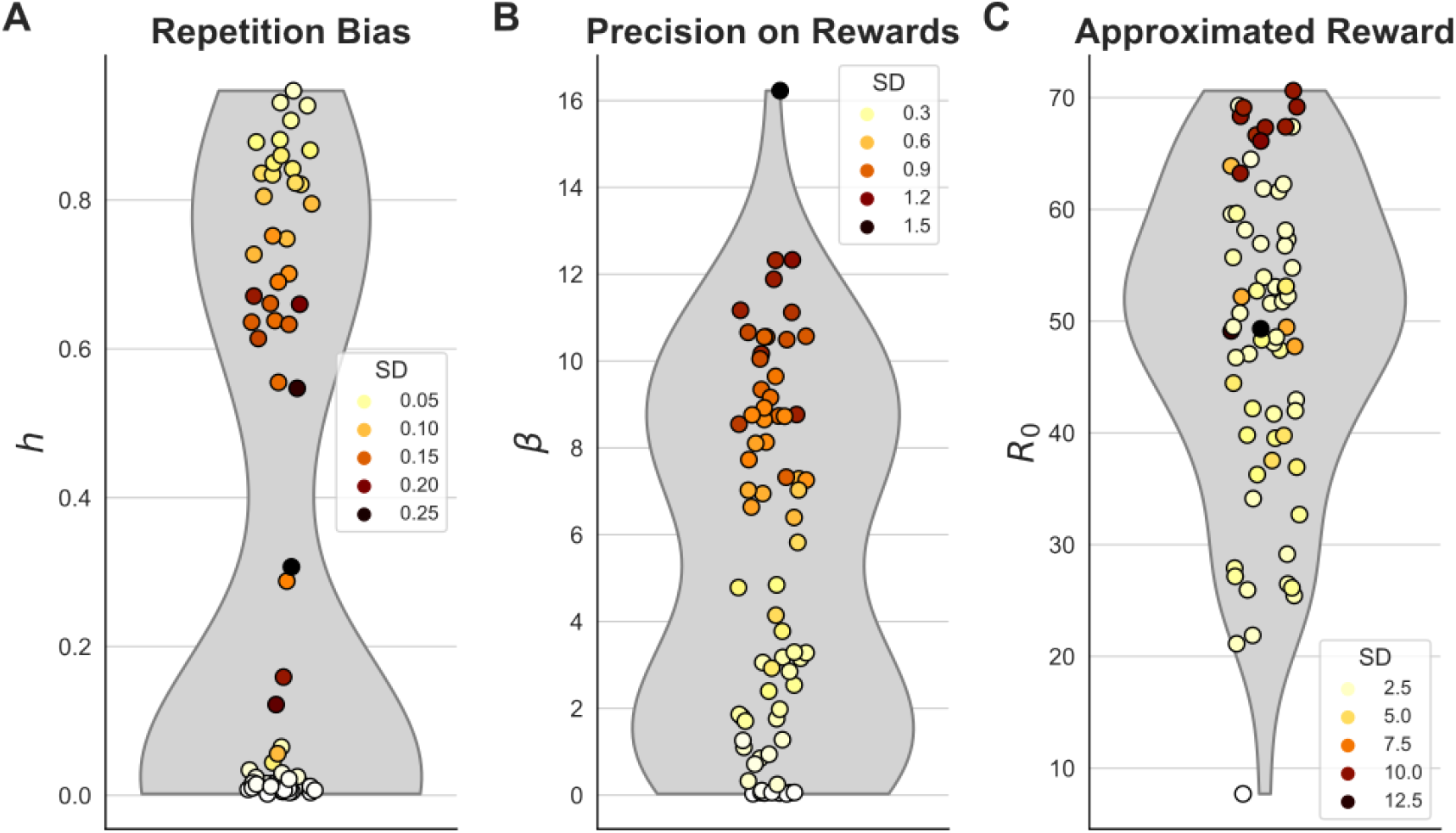
Estimated parameters of expected value with proxy and repetition bias model (EVPRM) Dots represent posterior means of individual parameter estimates. Each plot represents one of the three free parameters: **(A)** repetition bias *h*, **(B)** precision on expected rewards *β* and **(C)** approximated reward *R*_0_. Grey patches represent kernel density estimates. The color of dots indicate standard deviations (*SD*).

### Posterior predictive checks for EVPRM

we conducted posterior predictive checks (***Gelman et al., 2013***) to assess if the fitted EVPRM can replicate the behavior of the participants. We used the method from PyMC that simulates choices of 1,000 agents for each participant based on the model and posterior. The parameters of the agents were drawn from the posterior distributions.

We calculated the proportion of correctly predicted choices for each participant over all agents. These proportions of correctly predicted choices showed a very wide range from 4.9% to 86.3%, but all proportion were above the chance level of 2.7% (see ***Figure 11***A). On the group level the EVPRM predicted DAS choices better (74.9%) compared to non-DAS choices (19.1%, see ***Figure 11***C).

**Figure 11.**
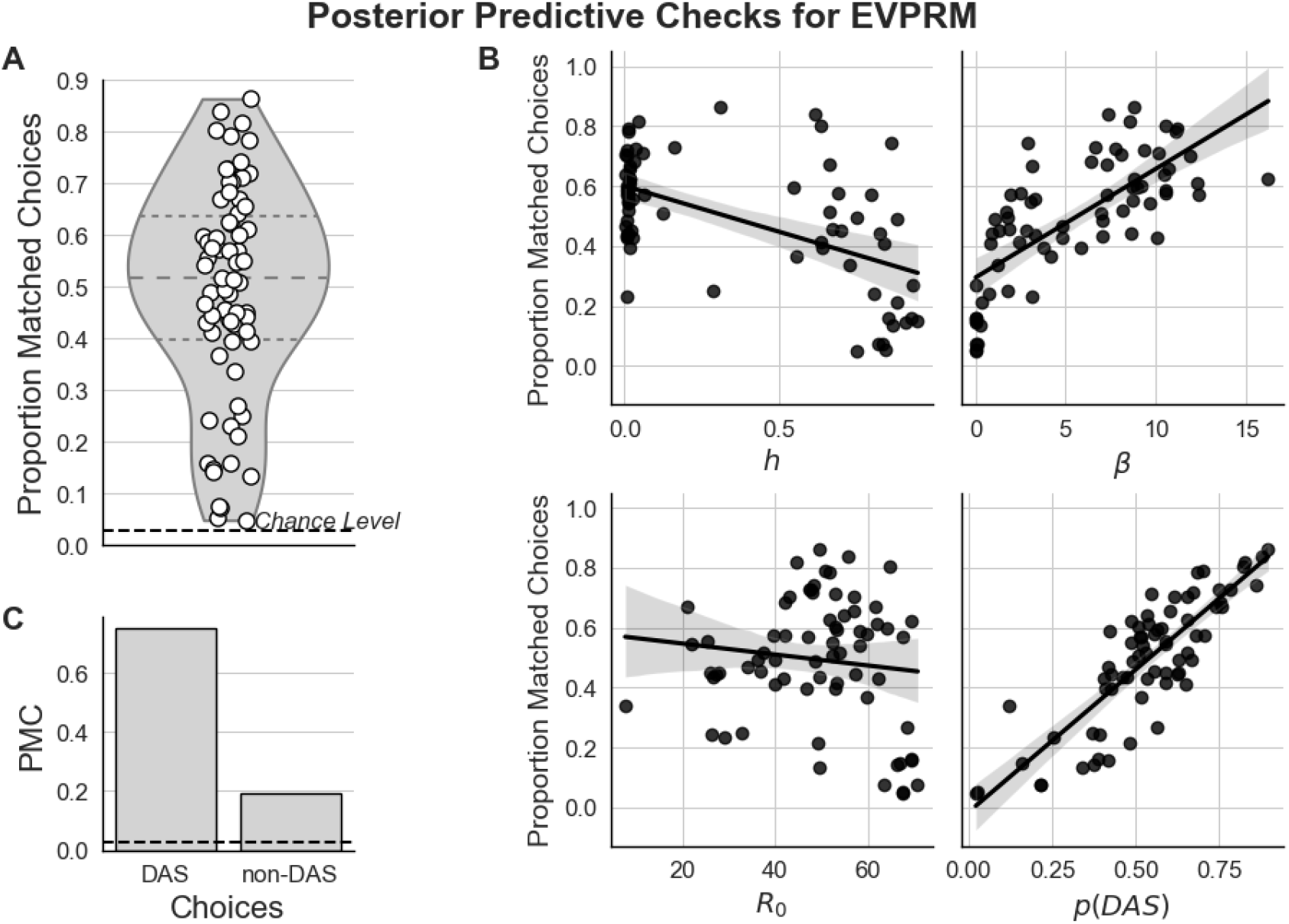
Posterior predictive checks (PPC) for expected value with proxy and repetition bias model (EVPRM). **(A)** Distribution of the proportion correctly predicted choices for each participant based on simulated data with the inferred parameters from the EVPRM. Each white dot represents the proportion of correctly predicted choices for one participant. The grey area represents a KDE of the distribution, and the dotted lines inside the KDE represent the borders of the quartiles. **(B)** Correlations between proportion of correctly predicted choices of each participant and their inferred parameters of the EVPRM and their proportion of default action sequence (DAS) choices. Black solid lines represent linear regression model fitted to the data. *h*: repetition bias, *β*: precision over expected rewards, *R*_0_: approximated reward, *p*(DAS): proportion of DAS choices. **(C)** Proportions of correctly predicted DAS and non-DAS choices at the group-level. Dotted line represents chance level. PMS: proportion matched choices.

We further calculated correlations between the proportions of matched choices and the posterior means of the inferred parameters and the proportion of DAS choices *p*(DAS). Here EVPRM better predicted choices of participants with weak repetition bias *h, r* = .55, *p <* .001, higher precision over expected rewards *β, r* = .73, *p <* .001, and higher proportions of *p*(DAS), *r* = .84, *p <* .001 (see ***Figure 11***B). The correlation with the approximated reward *R*_0_ was not significant, *r* = .12, *p* = .25.

## Funding

This work was funded by the German Research Foundation (DFG, Deutsche Forschungsgemeinschaft), SFB 940—Project number 178833530, and TRR 265—Project number 402170461 and as part of Germany’s Excellence Strategy—EXC 2050/1—Project number 390696704—Cluster of Excellence, Centre for Tactile Internet with Human-in-the-Loop (CeTI) of TUD Dresden University of Technology.

## Appendix

**Appendix 0—table 1.**
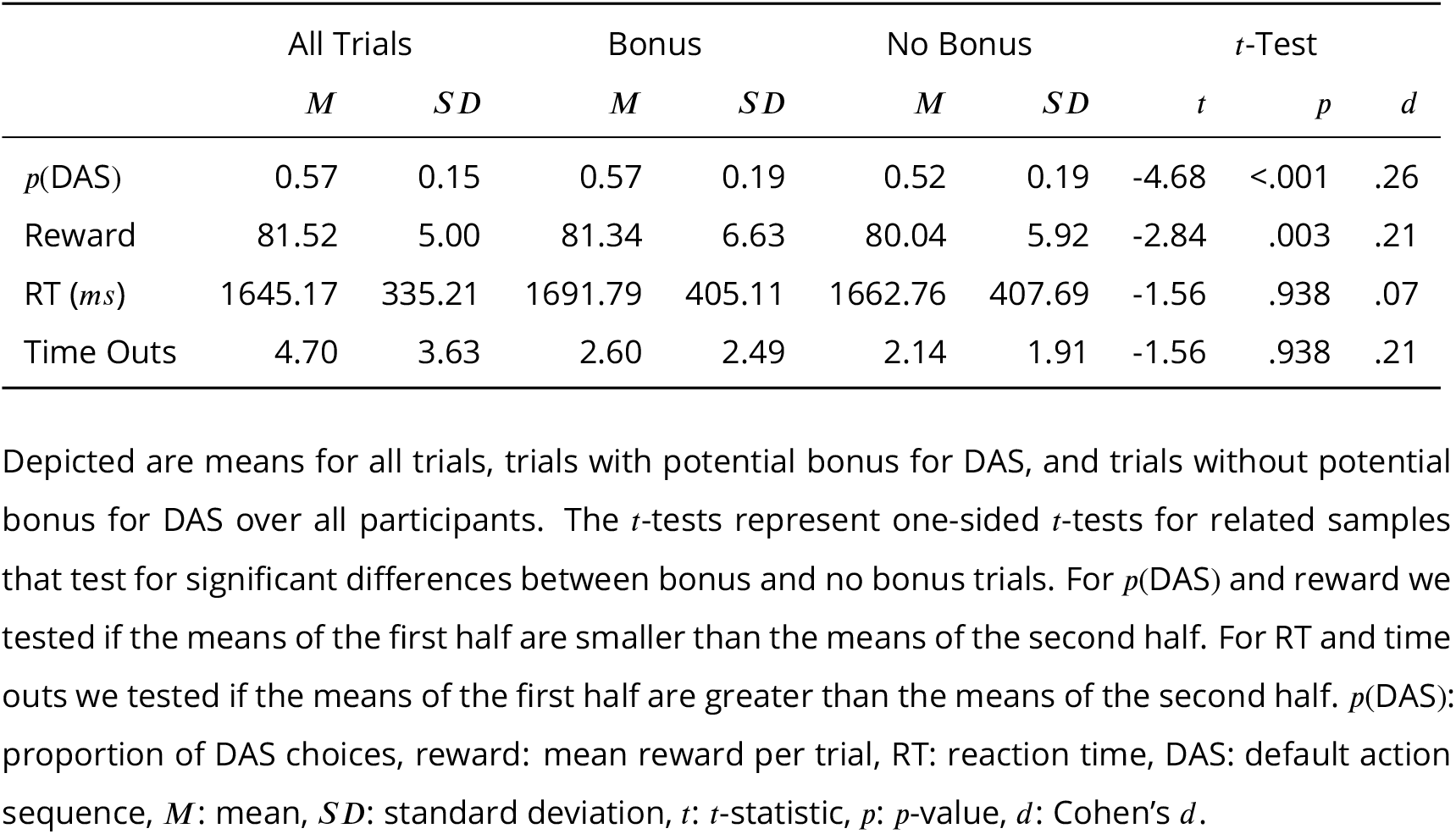
Descriptive Statistics depending on bonus condition

**Appendix 0—table 2.** Correlations between inferred parameter values of EVPRM and performance measures

| | $h$ | $\beta$ | $R_0$ | $p(\text{DAS})$ | Reward | RT | Time outs |
| --- | --- | --- | --- | --- | --- | --- | --- |
| $h$ | - | | | | | | |
| $\beta$ | -0.75 | - | | | | | |
| $R_0$ | **0.12** | **0.30** | - | | | | |
| $p(\text{DAS})$ | -0.22 | **0.37** | -0.18 | - | | | |
| Reward | **-0.69** | **0.76** | -0.11 | **0.67** | - |  |  |
| RT | **0.37** | **-0.44** | **0.25** | **-0.58** | **-0.70** | - |  |
| Time outs | **0.32** | **-0.27** | 0.16 | **-0.40** | **-0.52** | **0.65** | - |

Depicted are correlation coefficients between inferred parameter values of EVPRM and performance measures. As performance measures the means of each participant were used. Significant correlations are bold ( $p < .05$ ). $\beta$ : precision over expected rewards, $R_0$ : approximated reward, $h$ : repetition bias, $p(\text{DAS})$ : proportion of DAS choices, reward: mean reward per trial, RT: reaction time, DAS: default action sequence, EVPRM: expected value with proxy and repetition bias model.

**Appendix 0—figure 1.**
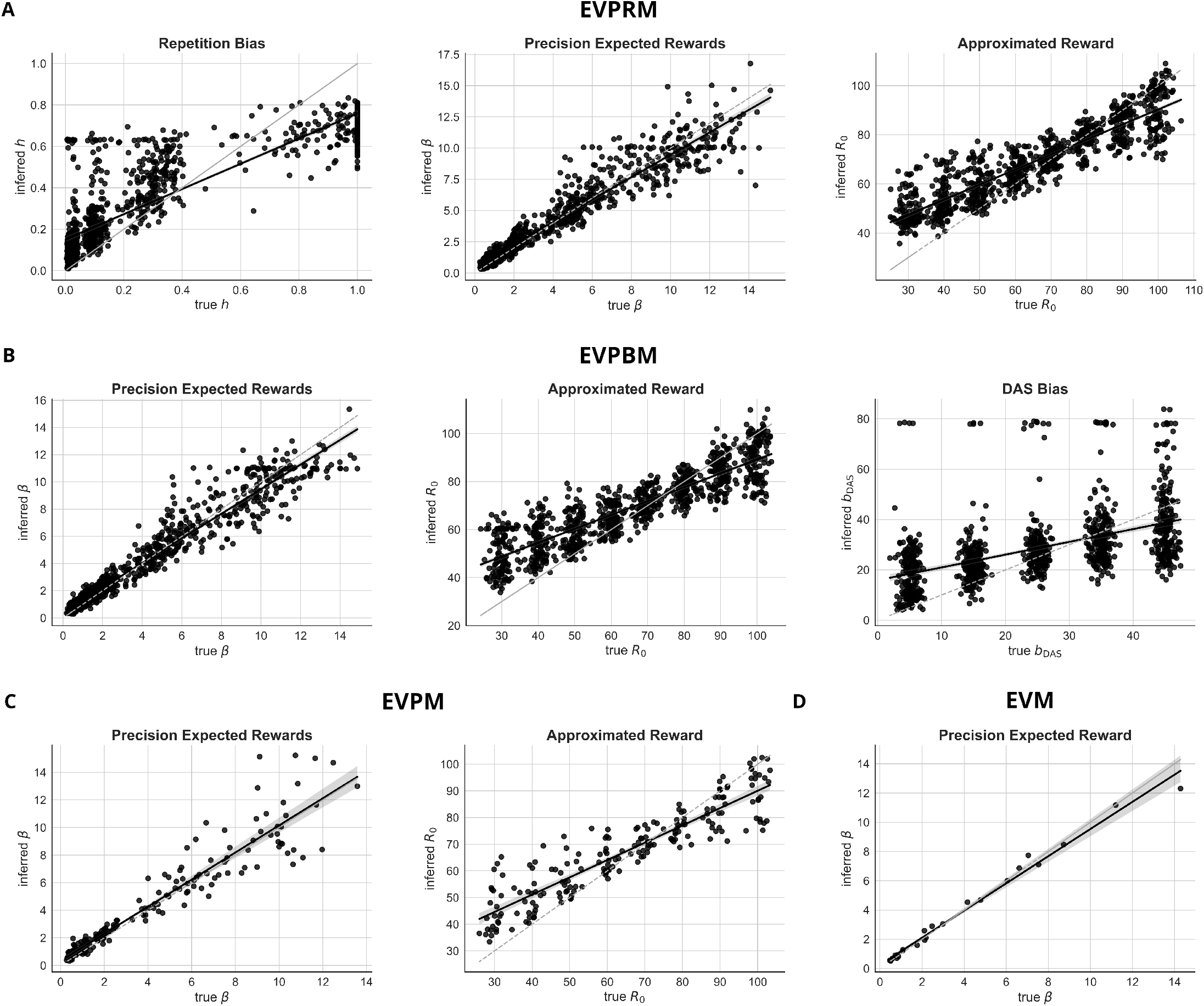
Parameter recovery for all candidate models. Correlations of true and inferred parameter values for all free parameters of the four candidate models: (**A**) expected value with proxy and repetition bias model (EVPRM), (**B**) expected value with proxy and default bias model (EVPBM (**C**) expected value with proxy model (EVPM), (**D**) and expected value model (EVM). Black solid lines represent correlation between true and inferred parameter values. Grey dashed lines represent true parameter values.

## Notes

https://github.com/ericleg/RepBias

### Competing Interest Statement

The authors have declared no competing interest.

